# Joint Analysis of Common Regulatory and Rare Coding Variants for Improved Power in Kernel Association Tests

**DOI:** 10.64898/2026.11.11.765949

**Authors:** Yuchen Yang, Eden R. Martin, Lindsay A. Farrer, Margaret A. Pericak-Vance, Jeffery M. Vance, Jonathan L. Haines, William S. Bush

**Author notes:** Contributing authors.

## Abstract

Rare alleles arise from mutations on an existing haplotype background, yet current gene-based rare variant association tests treat low-frequency coding variants as effects independent of the common regulatory variants that surround them. The penetrance of rare coding variants can be modulated by the cis-regulatory landscape, and failing to account for this context limits the statistical power of existing methods for complex traits. We present PrediXSKAT, a kernel-based rare variant association framework that incorporates the surrounding cis-regulatory landscape by estimating its impact on gene expression via PrediXcan. PrediXSKAT embeds predicted gene expression directly into the Sequence Kernel Association Test kernel through element-wise similarity weighting, concentrating rare variant signal among individuals who share concordant genetic and regulatory backgrounds. Comprehensive simulations demonstrate that PrediXSKAT increases power over standard kernel-based testing under biologically realistic conditions where a proportion of tested variants carry small, mixed-direction effects. Type I error rates are controlled in the absence of association. Power gains persist across a range of sample sizes and causal variant architectures. Applied to whole-genome sequencing data from the Alzheimer’s Disease Sequencing Project, PrediXSKAT recapitulates established disease loci and amplifies signals at previously unreported candidate genes without indiscriminately inflating genome-wide results. PrediXSKAT provides a generalizable and extensible framework for the joint analysis of common regulatory and rare coding variants. The approach can be applied to any kernel-based set test and extended to jointly consider multiple genes within the same biological pathway, offering a principled strategy for improving rare variant discovery in large-scale sequencing studies of complex disease.

## 1 Introduction

The availability of whole-genome sequencing in recent years has revealed a vast number of genetic variants in the human population. The landscape of human genetic variation is dominated by rare variants (RVs), typically defined as having a minor allele frequency (MAF) of less than 1%. For example, 97% of biallelic variants found through whole-genome sequencing in the Alzheimer’s Disease Sequencing Project (ADSP) Release 4 (n=45365) are rare, and 53% of variants are singletons, detected only in a single individual in the entire dataset [1]. Traditional single-variant association tests in genome-wide association studies (GWAS) are limited when applied to rare variants because the per-variant statistical power to detect associations decreases sharply with decreasing allele frequency. This challenge is compounded by the multiple testing burden inherent in genome-wide analyses, where millions of variants are tested simultaneously [2].

Variant set tests have been developed to estimate the joint effects of a group of variants, such as coding variants within a gene, and a phenotype. These approaches aggregate genetic signals within a biologically meaningful unit, thereby increasing statistical power by reducing the multiple testing burden. Leveraging the assumption that variants within the same functional unit may have similar effects on the phenotype, burden tests assume that all variants in a set act in the same direction and combine genotypes into a single score [3]. Alternatively, the Sequence Kernel Association Test (SKAT) allows for variants to have heterogeneous effect directions and magnitudes through a variance-components framework [4]. More recent approaches, such as the variant-Set Test for Association using Annotation infoRmation (STAAR), use an ensemble of burden tests and SKAT while weighting the variants with causal probability based on functional annotations [5].

However, these tests still exhibit limited statistical power when applied to complex polygenic traits. The heterogeneity of effect sizes and directions within variant sets, combined with the sparsity of truly causal variants, causes current methods to miss genuine biological signals [6]. Additionally, most existing methods treat all variants within a set equally or weight them solely based on allele frequency or predicted functional impact, without considering the broader genomic context in which these variants exist.

The penetrance of coding variants can be modulated by the cis-regulatory landscape through variation in gene and protein expression levels. For example, Castel et al. showed that pathogenic variants associated with cancer and autism were significantly enriched in patients with eQTL haplotype configurations correlating with higher expression [7]. In Alzheimer’s disease (AD), this has been explored through the lens of epistasis, where *APOE* (MIM:107741) is known to have additive and interaction effects with multiple other SNPs [8]. There is also evidence of interaction between high-effect *ABCA7* (MIM: 605414) loss-of-function variants and common moderate-risk variants to influence AD risk [9].

We hypothesize that incorporating regulatory variants with rare coding variants can improve RV association tests and reveal novel disease-associated genes. Our method extends the SKAT framework by weighting the rare variant kernel using gene expression levels predicted from surrounding expression quantitative trait loci (eQTLs) as a dosage for the functional gene product. This approach leverages population- and tissue-specific eQTL models to account for both population structure and trait-relevant regulatory variation, as gene regulation patterns differ substantially across ancestries and tissue types.

We term our method PrediXSKAT, as it integrates the PrediXcan framework for predicted gene expression with the SKAT rare variant test. PrediXcan is a widely used approach for estimating tissue-specific gene expression from genotype data using elastic net models trained on reference panels such as the Genotype-Tissue Expression (GTEx) project [10].

Results from extensive simulation studies show that PrediXSKAT provides improved statistical power for detecting signal from rare variant sets with sparse effect sizes and mixed effect directions, without inflating false positive rates for a range of sample sizes. We further apply PrediXSKAT to whole-genome sequencing data from the Alzheimer’s Disease Sequencing Project (ADSP) to identify known and previously unreported disease-associated genes. The ADSP represents one of the largest collections of whole-genome sequencing data for Alzheimer’s disease, providing an ideal testbed for evaluating the performance of our method on a complex neurodegenerative phenotype.

## 2 Methods

### 2.1 Whole-genome sequencing data

We analyzed Release 4 of whole-genome sequencing (WGS) data from the Alzheimer’s Disease Sequencing Project (ADSP) consisting of 45365 individuals with clinically-adjudicated AD case-control status. Highly related individuals were excluded according to guidelines from the Genome Center for Alzheimer’s Disease (GCAD) working group. Three subgroups were extracted for population-specific analyses. Individuals who reported their race as White and their ethnicity as non-Hispanic were classified as non-Hispanic White (NHW). Individuals who indicated any reported race and Hispanic ethnicity were classified as Hispanic (HIS). Individuals who reported their ethnicity as non-Hispanic and their race as Black were classified as African American (AA). Population structure was assessed using principal component analysis on a set of common, linkage disequilibrium-pruned variants. We included sex, PCs 1–5 (calculated through PC-AiR), and dosage of APOE *ε*4 and *ε*2 alleles as covariates, as well as dummy-encoded covariates for sequencing center, sequencing platform, read length, and whether PCR amplification was used. Table 1 shows demographic information for each subgroup.

**Table 1.** Alzheimer’s Disease Sequencing Project Release 4: Demographic and clinical characteristics by racial/ethnic group.

|  |  | NHW (*N* = 8692) | AA (*N* = 4250) | HIS (*N* = 6364) |
|---|---|---|---|---|
| AD |  | 4750 (54.6%) | 1286 (30.3%) | 2497 (39.2%) |
| Sex: Female |  | 5167 (59.4%) | 3119 (73.4%) | 4349 (68.3%) |
| *APOE* *ε*4 allele count | 1 | 3264 (37.5%) | 1583 (37.2%) | 1622 (25.5%) |
|  | 2 | 703 (8.09%) | 277 (6.52%) | 233 (3.66%) |
| *APOE* *ε*2 allele count | 1 | 678 (7.80%) | 734 (17.3%) | 605 (9.51%) |
|  | 2 | 23 (0.265%) | 50 (1.18%) | 10 (0.157%) |

% as percentage of race/ethnicity group

### 2.2 Simulation studies

To evaluate the Type I error rates and statistical power of our method, we performed simulation studies under several configurations. We selected the *ABCA7* gene because it represents a known Alzheimer’s disease susceptibility gene with empirically observed allele frequency distribution, local linkage disequilibrium structures, and allelic heterogeneity comparable to those seen across the exome in population sequencing studies. This choice allows us to simulate under genotype configurations that reflect realistic mutational and regulatory architectures, rather than relying on synthetic coalescent models. Figure 1 shows the structure of the *ABCA7* gene, annotated with loci of rare variants and eQTLs.

**Fig. 1.**
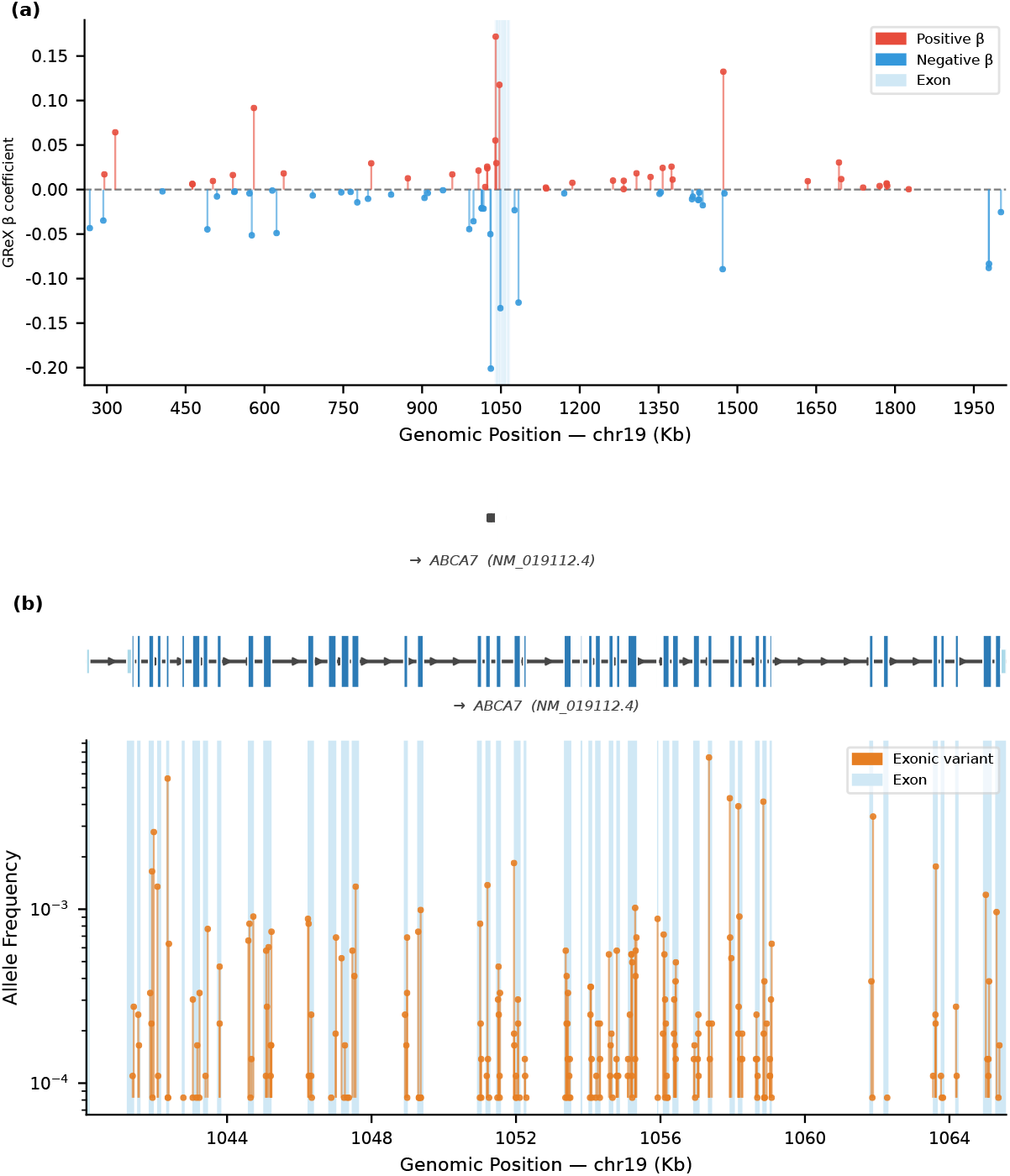
Regulatory and rare coding variant landscape at the ABCA7 locus. (a) Regulatory variants contributing to genetically regulated expression (GReX) of *ABCA7* in the GTEx(v8) whole-blood elastic net model. The y-axis shows the magnitude of the *β* coefficient for each variant; red and blue points denote variants with positive and negative *β*, respectively. The x-axis shows genomic position along chromosome 19; the shaded band marks the ABCA7 exon-intron structure, visible here only as a single block at this resolution. (b) Rare missense variants (MAF *≤* 1%) across ABCA7 exons, shown at higher genomic resolution. The y-axis shows allele frequency (log10 scale); shaded bands indicate individual exons resolved at this scale.

From the full ADSP dataset, we subsampled 1000 to 5000 individuals for various simulation scenarios. We used 162 missense coding variants within *ABCA7* with MAF*≤*1% and minor allele count MAC*>*2.

We simulated dichotomous traits from a logistic regression model to generate case-control phenotypes similar to Alzheimer’s disease. The baseline model without genetically regulated expression (GReX) effects was defined as

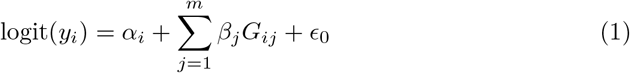

where *y_i_* denotes the case-control status for individual *i*, *α_i_* is a randomly generated covariate, *G_ij_* represents the genotype at variant *j*, and *β_j_* is the corresponding log odds ratio. The intercept term *ɛ*_0_ was adjusted to control overall trait prevalence.

Causal variant effect sizes were modeled as *β_j_* = *c|* log_10_ MAF*_j_|*, where *c* is set to 0.255 for most simulations, such that a variant with MAF = 5 *×* 10*^−^*^5^ has odds ratio OR = 3, and a variant with MAF = 0.01 has OR = 1.65. This parameterization reflects the empirically observed inverse relationship between allele frequency and effect size arising from purifying selection, whereby variants with larger effects tend to remain rare in the population.

We can then add a GReX term to our logistic models as an additive effect to rare variants, or also an interaction term with accumulated rare variants

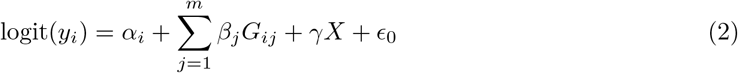

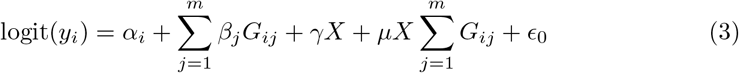

We also simulate a range of values for this coefficient *γ* that controls the effect size of GReX in the model.

To model heterogeneity within variant sets, we varied the proportions of variants with causal effects between 0%, 20%, 40%, 60%, 80%, and 100% by randomly selecting variants to set *β* = 0. The proportion of protective variants was varied across 0%, 10%, 20%, 30%, 40%, and 50% by randomly inverting the sign of selected effect sizes.

Each simulation configuration was repeated 200 times to obtain stable estimates of Type I error and statistical power. Power was computed as the proportion of replicates with p-values below significance thresholds: 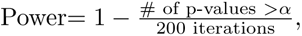 examined at *α* = 0.05, 0.005, 0.0005.

### 2.3 SKAT model and test

We use the simplest model of SKAT, formulated by Wu et al. 2011 [4].

We construct the kernel matrix **K** = **GWG**^⊺^ where **G** is an *n× m* genotype matrix of the region-of-interest with *n* individuals and *m* variants, where the (*i, j*)-th element is the genotype of variant *j* for individual *i*. **W** = diag(*w*_1_*, w*_2_*, … , w_m_*) is an *m × m* diagonal matrix where the diagonal elements 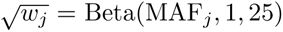 are weights for each variant on a beta distribution, such that rarer variants receive higher weights. We continue to use this weighting scheme due to its wide adoption even in newer formulations of SKAT.

The score test statistic is *Q* = (**y** *− π*)^⊺^**K**(**y** *− π*).

For our simulations, **y** is the vector of simulated case-control status. *π* is the vector of estimated case-control probability under the null hypothesis, calculated by fitting the null model logit(*y_i_*) = *α_i_* + *ɛ*_0_.

In real data analysis, **y** is the vector of clinically-adjudicated AD case-control status, while *π* is calculated by fitting the case-control status against the covariates using logistic regression without regularization.

We then use the following steps to estimate the null distribution for Q and obtain a p-value.

1. Define **V** = diag(*π*_1_(1 *− π*_1_)*, π*_2_(1 *− π*_2_)*, … , π_n_*(1 *− π_n_*)) where *π_i_* is the estimated case-control probability for the *i*-th individual under the null model.
2. Define **X̃** = [**1**, **X**] as a matrix of covariate columns concatenated with a column of ones.
3. Define **P** = **V** *−* **VX̃**^⊺^(**X̃**^⊺^**VX̃**)*^−^***^1^X̃**^⊺^**V**.
4. The eigenvalues of 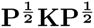 and the test statistic *Q* are then used to obtain a *p*-value from a null distribution estimated via Davies exact method [11]. We specifically use Davies method as implemented through FaST-LMM.

### 2.4 GReX kernel

For simulations, we used whole-blood eQTL elastic net models from GTEx(v8) with PrediXcan to compute predicted genetically regulated expression (GReX) [10]. We then use quantile transformation to a uniform distribution to assign each individual a value in [0, 1]. This transformation ensures that the GReX kernel has similar scale properties regardless of the specific gene being analyzed and prevents genes with very different expression ranges from dominating the kernel computation. The quantile transformation also makes the method robust to outliers and unusual distributions in predicted expression values.

Pairwise differences in transformed values are used to construct a kernel of GReX similarity between individuals *K*_GReX_. We base this on a radial basis function kernel, formulated as 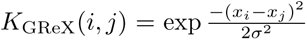 for individuals *i* and *j* with transformed GReX values *x_i_* and *x_j_*. *σ* is the median of non-zero absolute values of pairwise differences. We then scale this pairwise similarity measure between 0.5 and 2, such that individuals with identical predicted expression receive the maximum kernel value, while individuals with highly different expression receive a substantially reduced but non-zero kernel value. These specific parameterizations were selected through preliminary testing to ensure that the resulting modification of rare variant similarity is meaningful but not extreme.

We incorporate the GReX kernel into the SKAT test by forming a new kernel **K**_PrediXSKAT_ = **K_rare variants_** *⊙* **K**_GReX_, which is the element-wise product of the two kernel matrices. This operation up-weights the similarity of rare variant genotypes between two individuals if they also have similar predicted gene expression levels, and conversely down-weights rare variant similarity if the predicted gene expression levels are substantially different. The PrediXSKAT kernel is positive semidefinite and asymmetrically encodes joint similarity to amplify signal while bounding potential attenuation.

**K**_PrediXSKAT_ is then passed to SKAT for significance testing.

## 3 Results

### 3.1 Simulations

We evaluated the Type I error and power of PrediXSKAT compared to conventional SKAT using a simulated dichotomous trait under a range of configurations. Type I error rate was evaluated based on 200 iterations at *α* = 0.05, 0.005, 0.0005. We estimated power as the proportion of p-values less than the set *α* level based on 200 replicates in each condition. We considered different proportions of causal variants and mixtures of effect directions in the variant set.

#### 3.1.1 Simulated case-control baseline

For the simplest model where only the rare variants have an effect on the simulated phenotype, we use 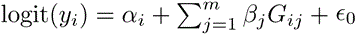 where for each individual *i*, covariate *α_i_ ∼ N* (0, 1) is randomly generated, *β_j_* is the log odds ratio (OR) of the *j*-th variant, and *G_ij_* represents the genotype for the *m* rare variants in the region. The intercept term *ɛ*_0_ was calibrated to create a trait prevalence of 0.46, matching the case-control ratio in the full ADSP dataset.

Simulation results are shown in Figure 2. Across all scenarios, PrediXSKAT consistently outperforms the unmodified SKAT. Including GReX as a covariate does not improve power consistently. Under the null model (no causal variants), power approaches zero, confirming well-controlled Type I error rates (Figure S1). When all causal variants act in the same direction, both methods achieve high power at higher causal proportions, but PrediXSKAT maintains superior performance even when only 50% of variants contribute to disease risk.

**Fig. 2.**
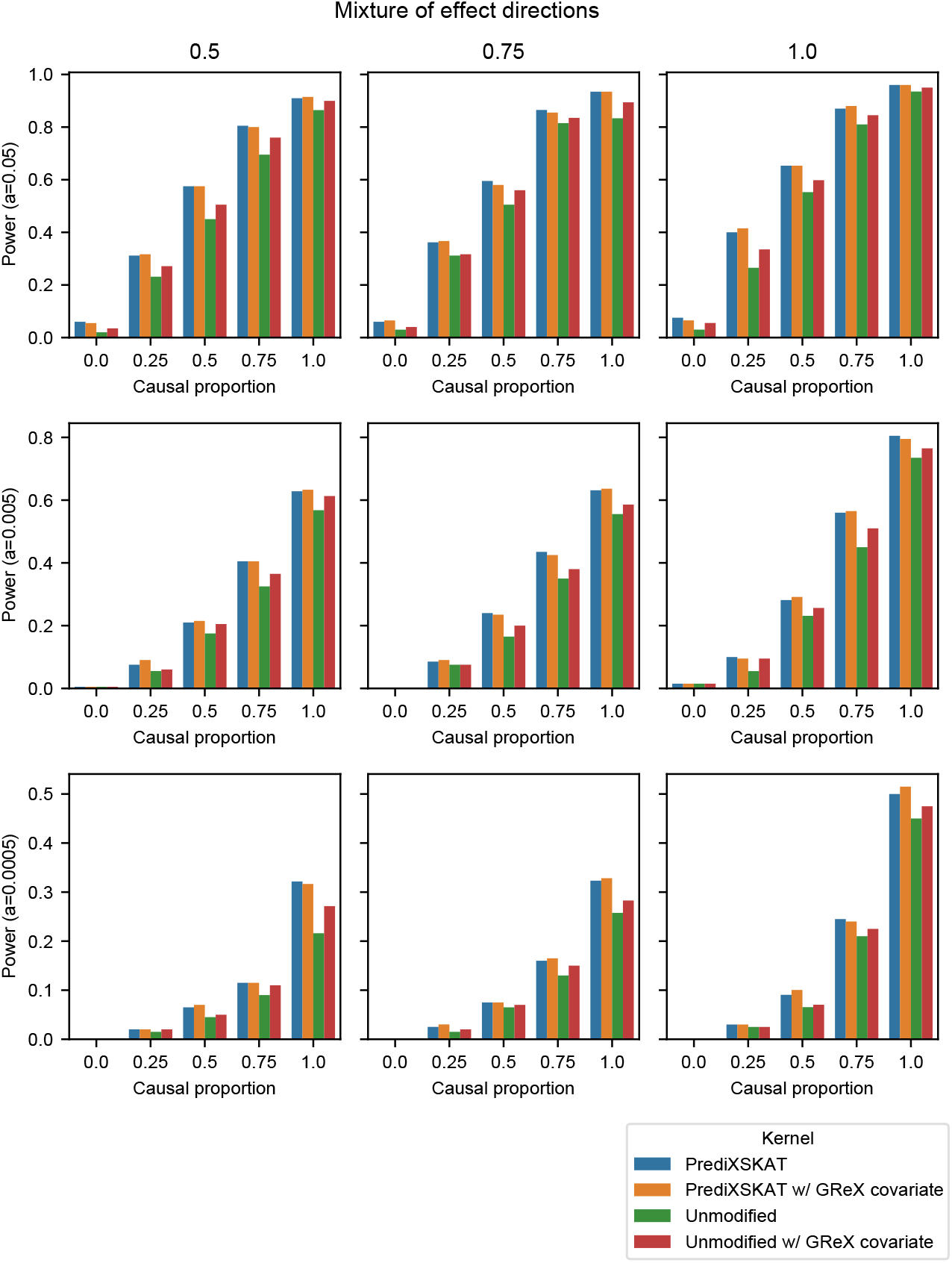
Baseline power of PrediXSKAT versus unmodified SKAT for a simulated case-control trait. Columns correspond to increasing proportions of deleterious vs protective variants (0.5, 0.75, 1.0). Rows show power at *α* = 0.05, 0.005, and 0.0005. Within each panel, the x-axis gives the proportion of rare variants with a non-zero effect size, and the y-axis gives empirical power calculated as the percentage of 200 simulation replicates (*n* = 2000 individuals) with p*< α*. “Unmodified” is the standard SKAT test without GReX. “PrediXSKAT” incorporates GReX into the kernel. Comparing kernels with and without GReX additionally included as a covariate tests whether a marginal effect of expression on phenotype, rather than the kernel modification itself, explains any power gain.

#### 3.1.2 Simulated case-control with additive GReX term

Phenotypes were simulated according to 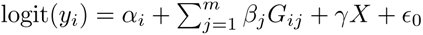 using *γ* = 0.1. Across all simulation settings, PrediXSKAT maintains well-controlled Type I error rates while consistently improving statistical power relative to SKAT (Figure 3).

**Fig. 3.**
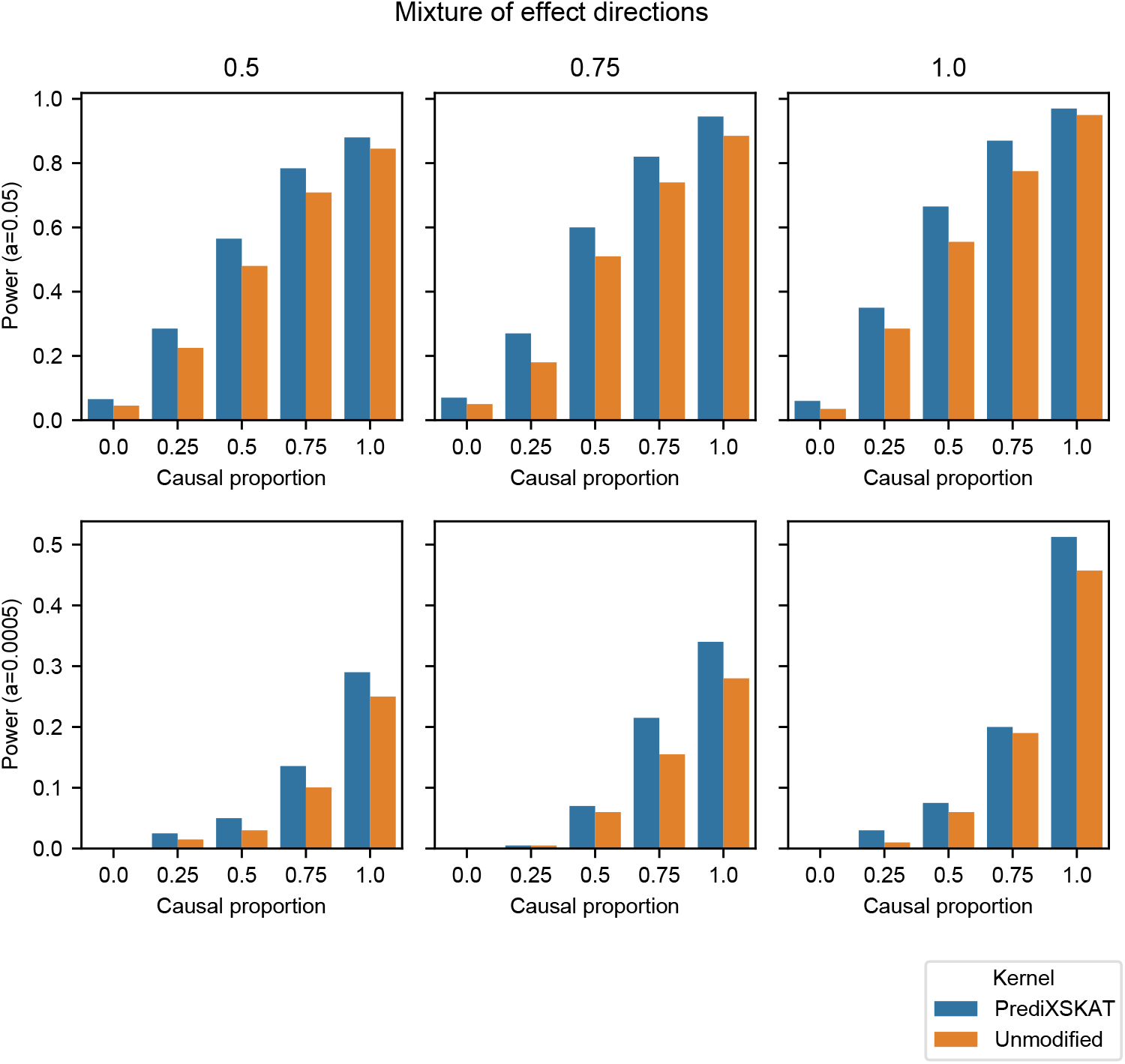
Power under a case-control model with an additive GReX effect on the simulated trait. Empirical power of PrediXSKAT versus unmodified SKAT, plotted as in Figure 2, across the same range of causal-variant proportions and effect-direction mixtures.

Increasing the GReX term effect size to *γ* = 0.5 results in a larger improvement in statistical power, as the aggregate effect of rare variants is reduced relative to the effect of predicted gene expression (Figure 4).

**Fig. 4.**
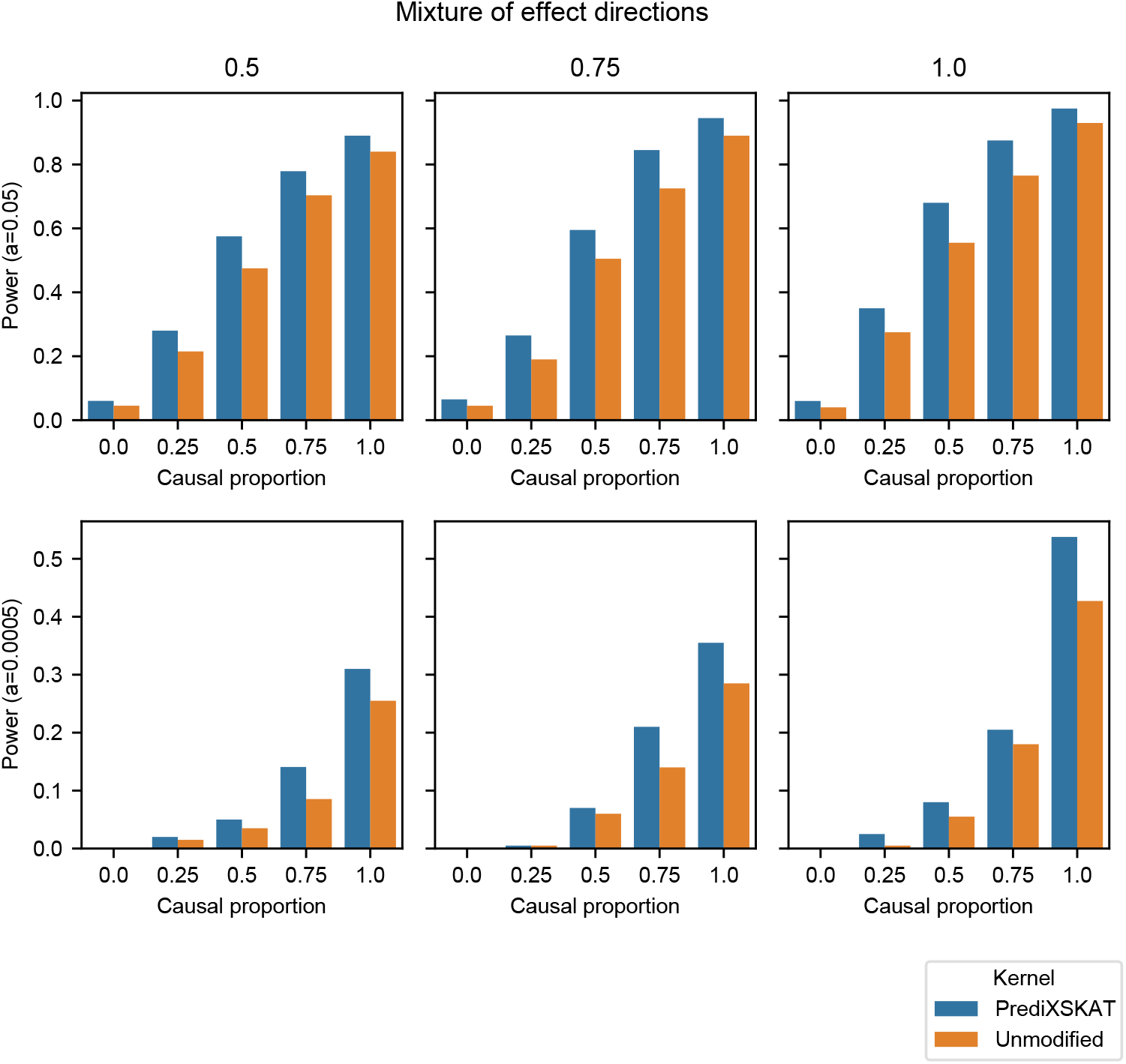
Power under a case-control model with a larger additive GReX effect (*γ* = 0.5; **Equation 2).** Empirical power of PrediXSKAT versus unmodified SKAT, plotted as in Figure 2, using the same simulation and replicate scheme (200 replicates, n = 2,000).

#### 3.1.3 Simulated case-control with additive GReX term and interaction term

We further tested a more complex model that included both a main effect of GReX and an interaction between GReX and rare variant burden: 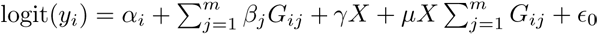 where *γ* = 0.1 and *µ* = 0.2. This model represents scenarios where gene expression not only has a direct effect on phenotype but also modulates the magnitude of rare variant effects. We found similar performance and improvements over SKAT, which is expected since the genotype matrix for rare variants is often sparse, leading to the simulated epistatic effect being also be proportionately small (Figure 5).

**Fig. 5.**
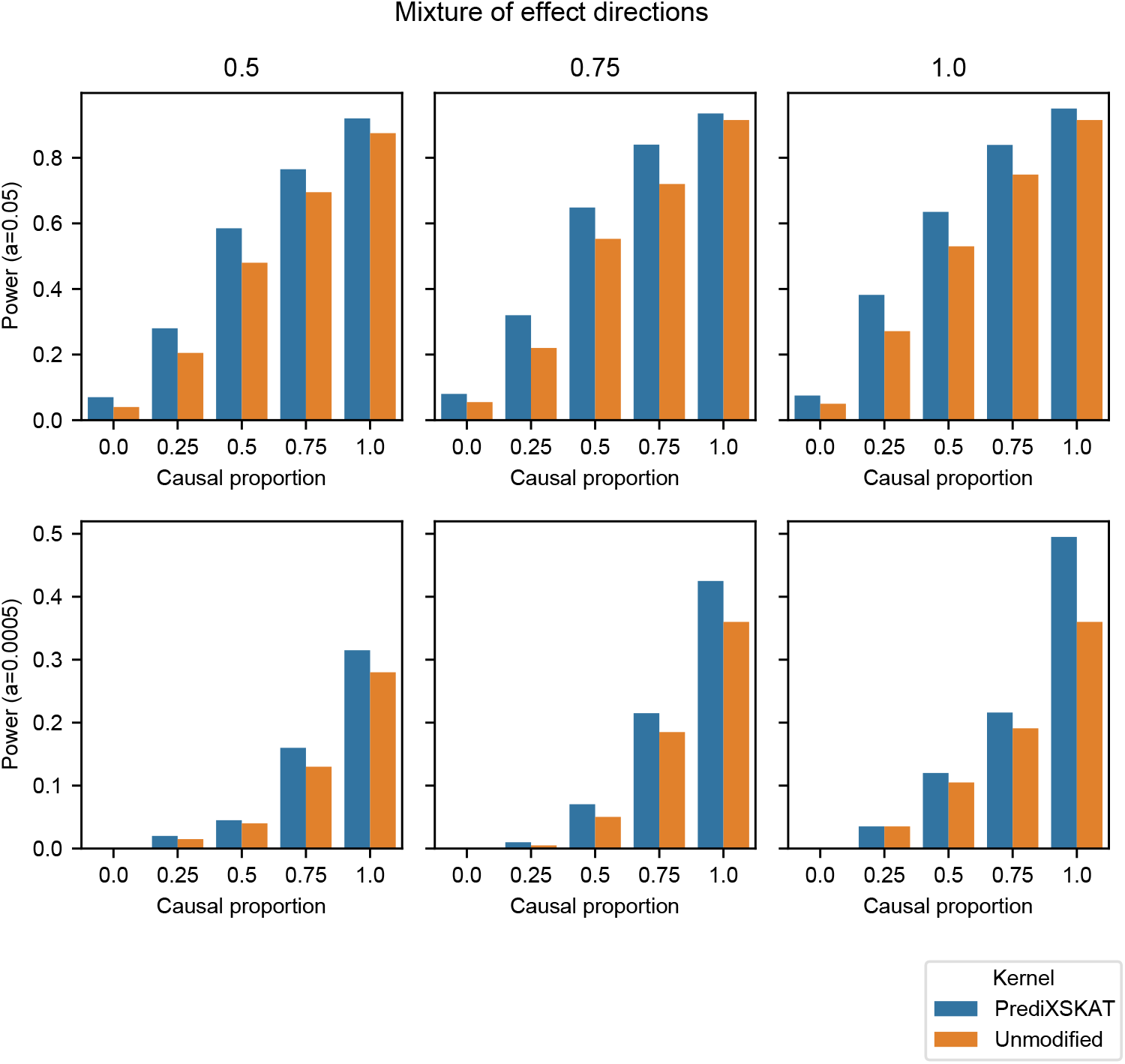
Power under a case-control model with additive and interaction GReX effects (*γ* = 0.1, *µ* = 0.2; **Equation 3).** Empirical power of PrediXSKAT versus unmodified SKAT, plotted as in Figure 2. GReX is included as a covariate in the trait-simulation model in addition to its role in the PrediXSKAT kernel.

#### 3.1.4 Simulating different effect sizes and different sample sizes under the additive GReX model

In Figure 6, we assessed the power of PrediXSKAT under a range of sample sizes (*n* = 2000, 3000, 4000, 5000) and different effect sizes for the GReX component (*γ* ranging from 0 to 5.0). These analyses provide practical guidance for study design and help identify scenarios where PrediXSKAT provides the greatest benefit.

**Fig. 6.**
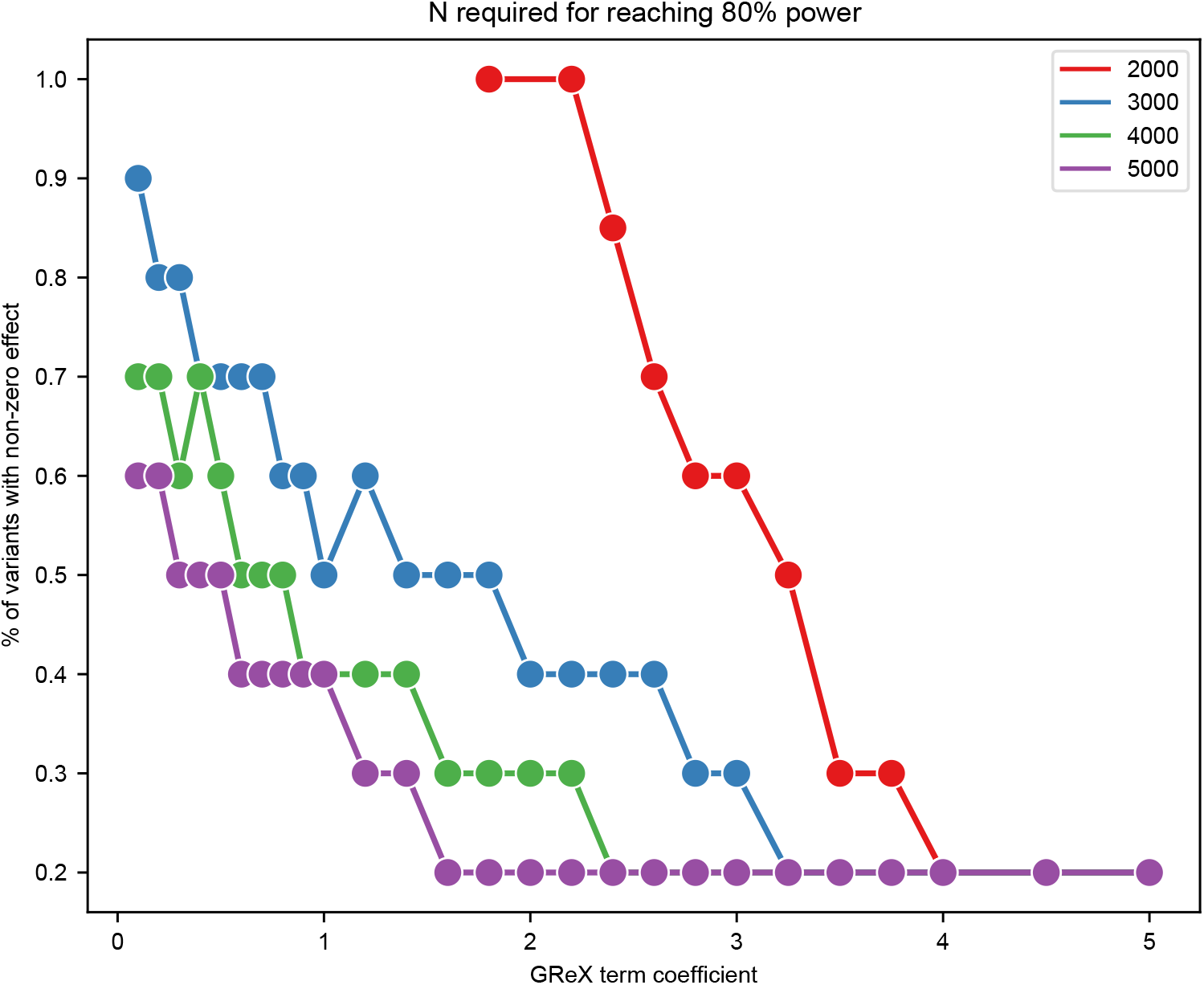
Minimum conditions for reaching 80% power using PrediXSKAT. Each line shows, for a given sample size (n=2000 to n=5000), the combinations of the GReX effect size (*γ*, x-axis) and the proportion of rare variants with a non-zero effect (y-axis) at which power first reaches 80% (*α* = 0.05). Simulations used a 50/50 mixture of protective and deleterious effect directions.

We tested a range of causal variant proportions and *γ* coefficients on the GReX term in the trait simulation. We use 50/50 mixed effect directions. For example, if a variant set-of-interest containing less than 30% variants with non-zero effect sizes, assuming a coefficient of 1.0 for GReX in the model, we would need more than 5000 case-control samples to reach 80% statistical power, assuming a similar prevalence. Alternatively, if more than half the variants in a set-of-interest have non-zero effects, and that gene expression has a strong effect as well corresponding to GReX larger than 2, we would only need between 2000 and 3000 samples to have 80% power in testing for these rare variants’ effects.

#### 3.1.5 P-value distributions under genome-wide significance thresholds

These simulations were done using an *α* between 0.05 and 0.0005. Real-world analysis with gene-based rare variant tests typically applies a Bonferroni-corrected threshold of approximately 2 *×* 10*^−^*^6^, when 20,000 protein-coding genes are tested. To understand the practical utility of PrediXSKAT under these stringent thresholds, we examined the full distribution of p-values across simulation replicates.

Figure 7 shows the distribution of p-values from the simulations using a model with additive GReX effect, where all variants in the tested set were given a non-zero effect in the same direction. A variant with MAF = 5 *×* 10*^−^*^5^ is set to OR=3, and a variant with MAF = 0.01 is set to OR=1.65. This represents an optimistic scenario where all variants in the set are causal, acting in the same deleterious direction, and having relatively strong effect sizes. Even in such an ideal scenario, we see that most p-values do not fall below threshold values typically used in the literature.

**Fig. 7.**
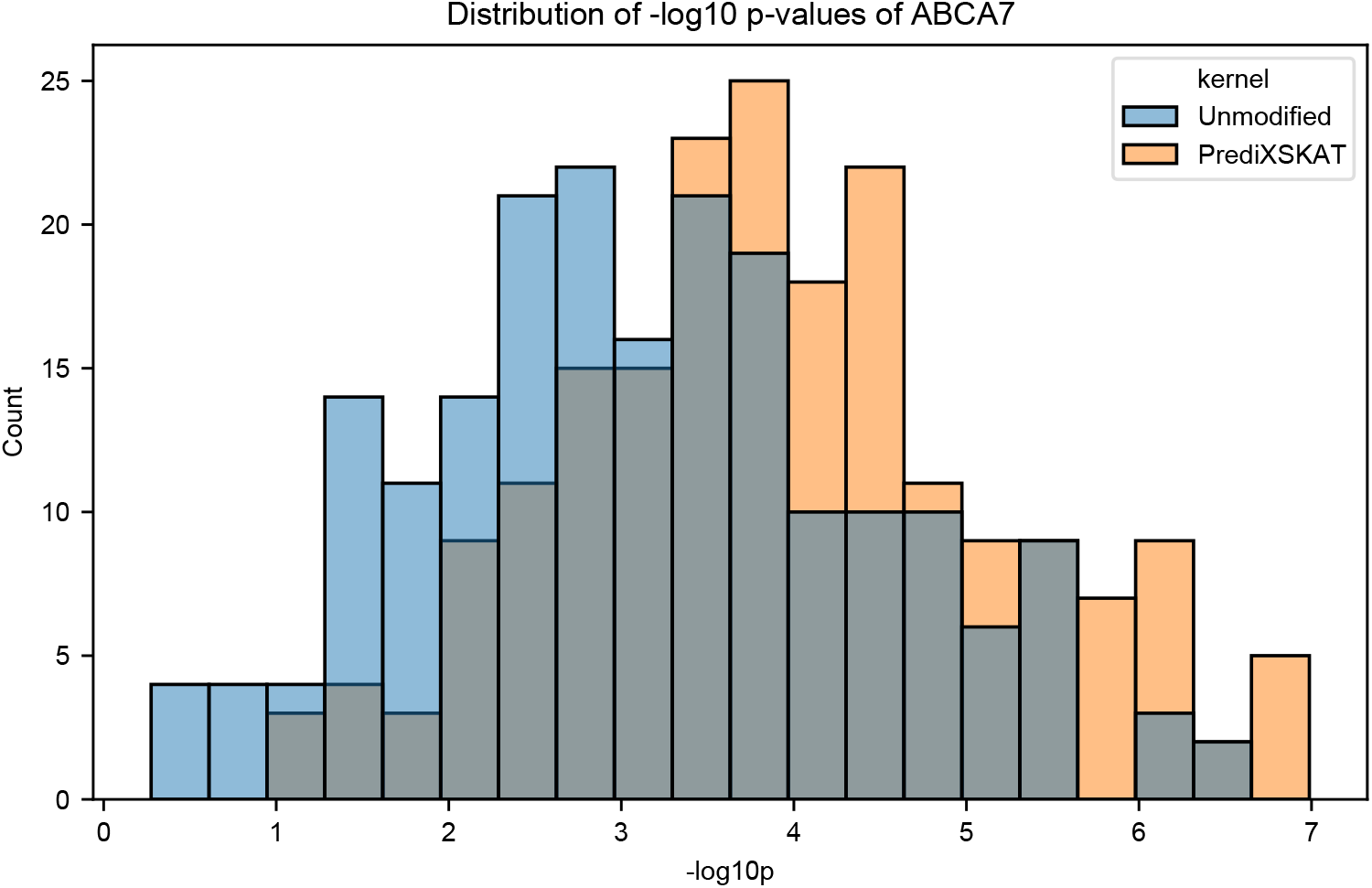
Distribution of -log10 p-values from simulations at the ABCA7 locus using a model with an additive GReX effect. All tested variants were assigned a non-zero deleterious effect scaled by minor allele frequency (a variant with MAF = 5 *×* 10*^−^*^5^ set to OR = 3; MAF = 0.01 set to OR = 1.65). The smallest observed p-value was 1.03 *×* 10*^−^*^7^.

### 3.2 Real data analysis

We applied our model to 3 subgroups of ADSP Release 4 (non-Hispanic White, African American, and Hispanic) to test exome-wide rare missense variants aggregated in gene-based sets. We restrict rare missense variants to those with MAC*>* 2 and MAF*<* 0.01 study-wide. GTEx(v8) whole-blood data, which was trained on NHW gene expression data (*n* = 670), was used to generate genetically regulated expression scores (GReX) with the PrediXcan software. We also used PrediXcan models of brain cortex and hippocampus. Hispanic and African American analyses were done using whole-blood GReX models from Kachuri et al. 2022 [12].

From Figure 8, we found that the majority of genes cluster close along the diagonal, indicating that for most genes PrediXSKAT and SKAT produce concordant p-values. This showed that PrediXSKAT does not indiscriminately inflate all signals. A subset of genes above the diagonal line represents genes where PrediXSKAT detected stronger signal than SKAT. Most of these genes already have a moderate SKAT signal, which is amplified by PrediXSKAT. The largest improvement was found in *PKHD1L1*, where the p-value was elevated from 6.60 *×* 10*^−^*^7^ to 1.91 *×* 10*^−^*^8^. Similar patterns were found when this was repeated using brain cortex and hippocampal GReX models (Figure S3).

**Fig. 8.**
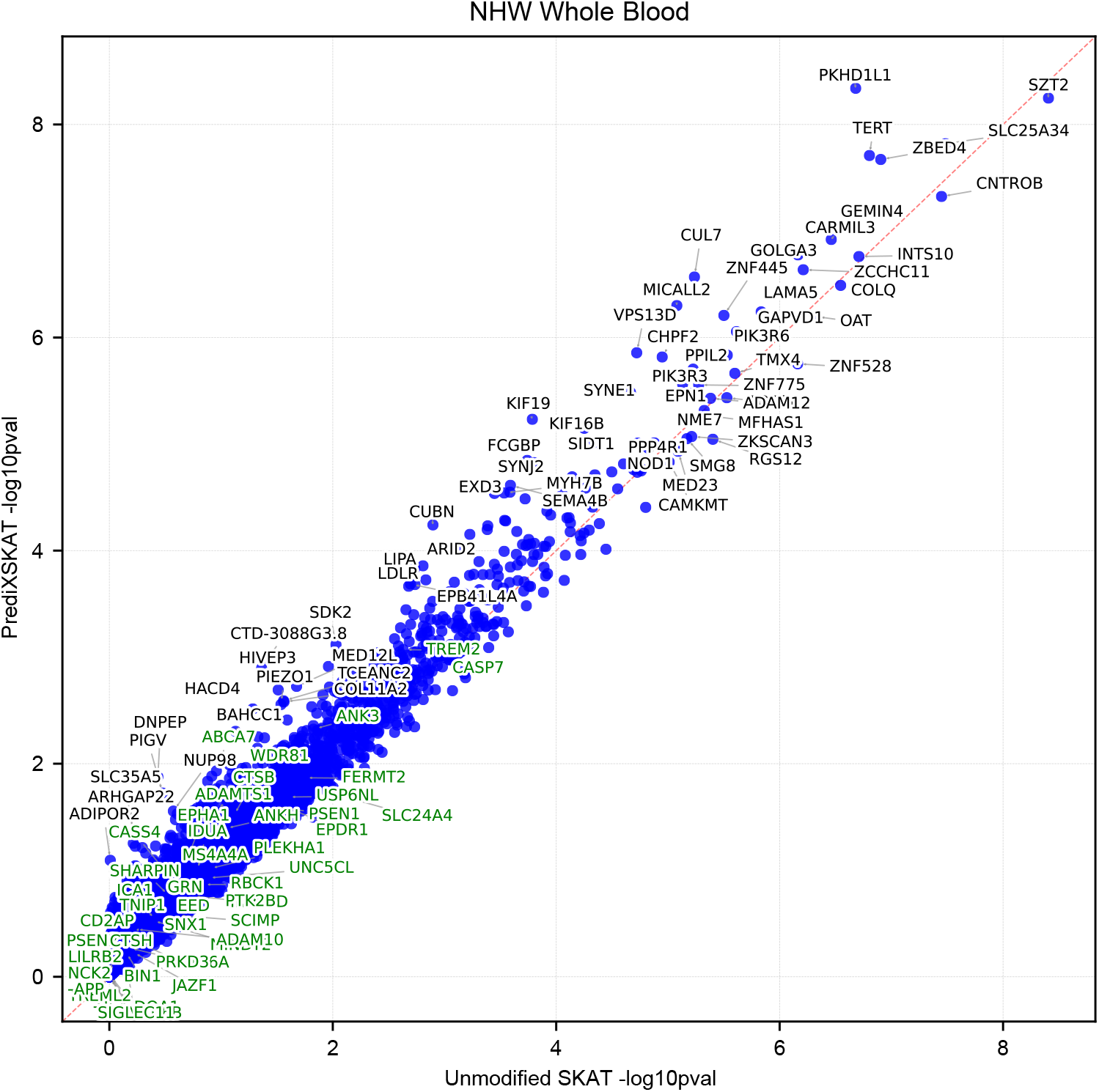
Gene-based rare-variant test results for the non-Hispanic White (NHW) subgroup using whole-blood GReX. Each point represents one gene tested. Axes show -log10-transformed p-values from unmodified SKAT (x-axis) and PrediXSKAT (y-axis). The dotted diagonal line marks identity. Genes labeled in green are previously reported AD loci verified by the ADSP committee. The 30 genes furthest from the diagonal, together with any gene at -log10p*>* 5, are labeled in black.

Plotting the distribution of p-value change from SKAT to PrediXSKAT shows that while most genes experience little change, genes that are affected by PrediXSKAT are more likely to show an increase in -log10p across the genome. The distribution cuts off around -0.5, meaning PrediXSKAT rarely and only modestly attenuates a strong SKAT signal (Figure 9). We also show Q-Q plots in Figure S2, where the genomic inflation factor (*λ*) increases from 1.198 with unmodified SKAT to 1.348 using PrediXSKAT.

**Fig. 9.**
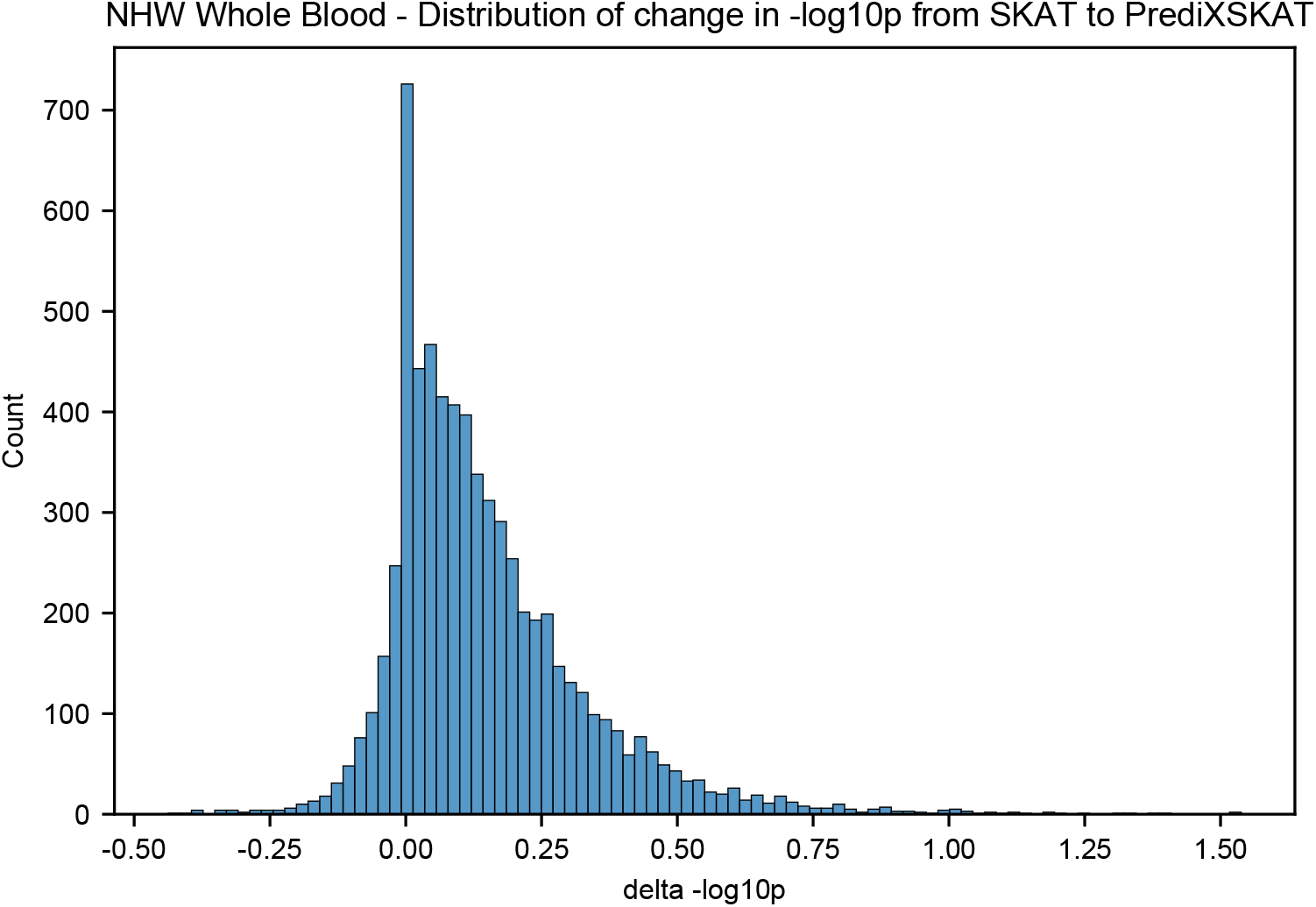
Distribution of the change in -log10 p (PrediXSKAT minus unmodified SKAT) across genes in the NHW whole-blood analysis shown in. **Figure 8**. Histogram of Δ-log10p between SKAT and PrediXSKAT. Positive values indicate a stronger PrediXSKAT signal relative to unmodified SKAT.

A cluster of genes (*SPTLC2* (MIM: 605713), *VASH1* (MIM: 609011), *ACOT4* (MIM: 614314), *PSEN1* (MIM: 104311)) on chromosome 14q24 has previously been reported to have significant associations in Hispanic individuals, and here we recapitulated those findings in the same ancestry group [13]. PrediXSKAT did not affect the signal of those genes (Figure 10).

**Fig. 10.**
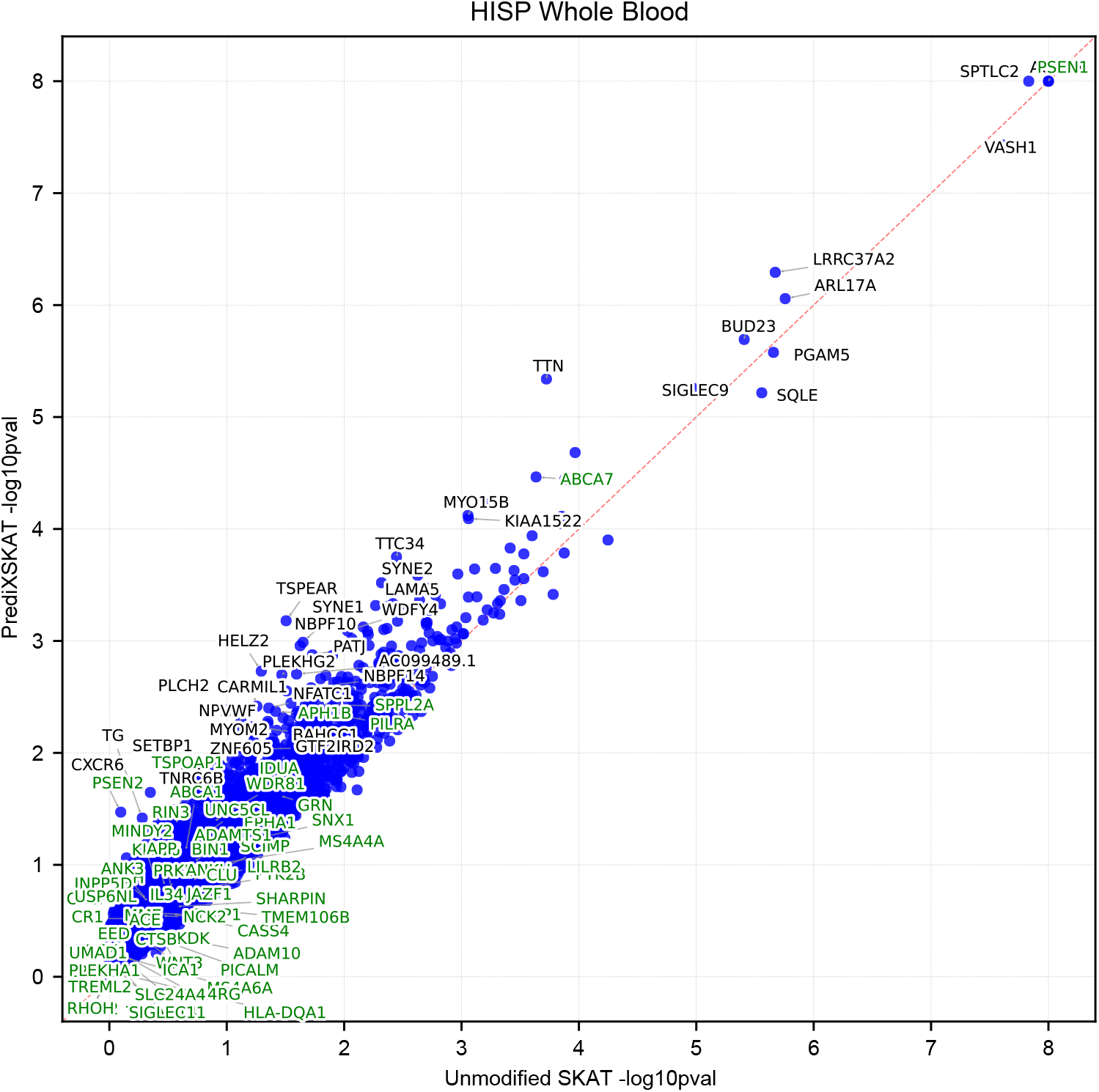
Gene-based rare-variant test results for the Hispanic (HIS) subgroup using whole-blood GReX. Plotted as in Figure 8. Genes labeled in green are previously reported AD loci verified by the ADSP committee. The 30 genes furthest from the diagonal, together with any gene at -log10p*>* 5, are labeled in black.

In the Hispanic subgroup results, we also see a second cluster of genes. *LRRC37A2* (MIM: 607843) and *ARL17A* are both on chr17q21.31, close to the *MAPT* (MIM: 157140) gene. This region is defined by a common inversion polymorphism generating the H1 and H2 haplotypes, with H1 associated with increased risk for tauopathies including Parkinson’s disease and progressive supranuclear palsy [14]. Fine-mapping studies of Parkinson’s disease risk at this locus have specifically identified *LRRC37A2* expression in astrocytes as a candidate mediating mechanism, with protective sub-haplotypes associated with increased *LRRC37A2* copy number and expression in human brain tissue. These two genes may be tagging the same *MAPT* regulatory signal instead of representing independent gene-level findings.

In the African American subgroup (Figure 11), no genes passed the Bonferroni-corrected significance threshold (*p <* 10*^−^*^5^). However, we observed PrediXSKAT having a larger amplification effect on more genes in AA than in the other ancestry groups. For example, the p-value of *EPPK1* (MIM: 607553) shifts 2 orders of magnitude from 0.03163 to 0.000204.

**Fig. 11.**
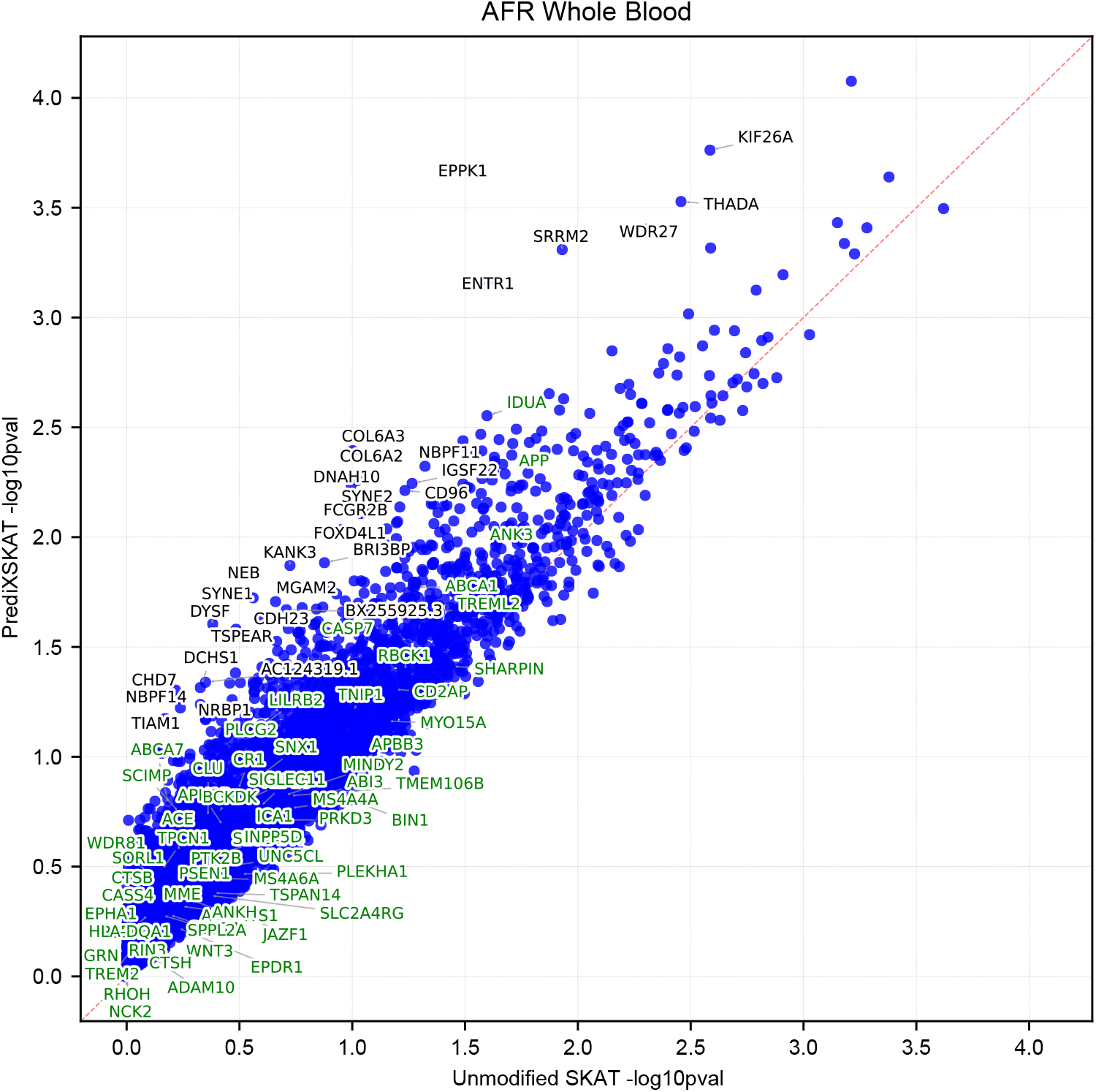
Gene-based rare-variant test results for the African American (AA) subgroup using whole-blood GReX. Plotted as in Figure 8 and 10.

We also hypothesized that we may see a more significant improvement if we restrict rare variants to those with high Phred-scaled Combined Annotation Dependent Depletion (CADD) score *≥* 20, as a way to filter out variants that likely have no or small effects, resulting in a variant set that has a high proportion of likely causal variants [15]. However, we found that this reduced the signal of PrediXSKAT in most genes (Figure 12).

**Fig. 12.**
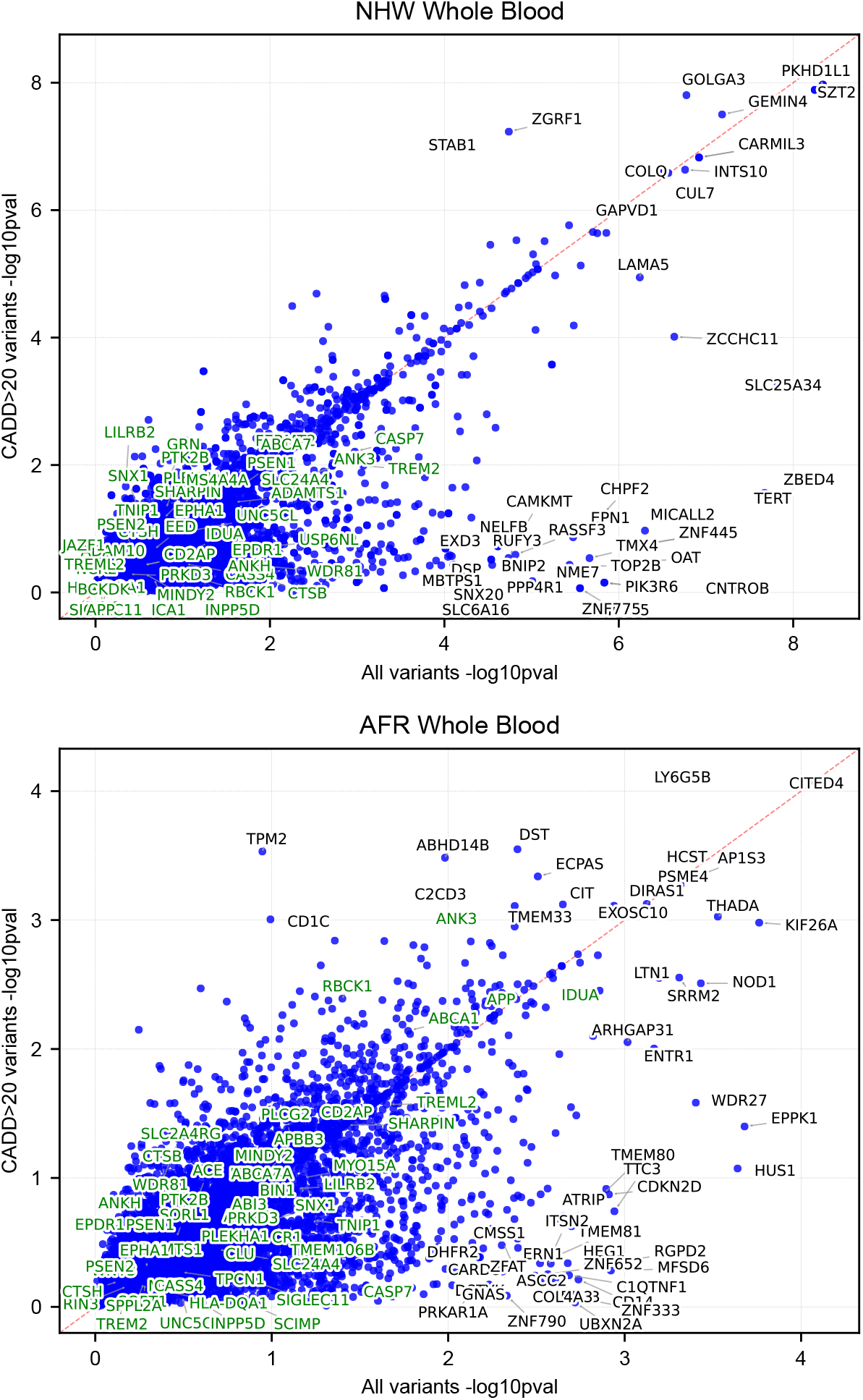
Effect of restricting to high-impact variants on the PrediXSKAT signal. (a) NHW whole-blood analysis. (b) AA whole-blood analysis. In each panel, the x-axis shows the PrediXSKAT -log10 p-value using all rare variants in a gene, and the y-axis shows the PrediXSKAT -log10 p-value using only variants with Phred-scaled CADD*≥* 20.

Overall, the PrediXSKAT analysis demonstrates that integrating predicted gene expression into rare variant testing can prioritize both established and previously unreported candidate genes. These findings suggest that accounting for genetically regulated expression in gene-based tests can reveal complementary aspects of disease biology not captured by variant-based models alone.

## 4 Discussion

In this study, we present PrediXSKAT, a gene-based rare variant association framework that integrates genetically regulated gene expression (GReX) into the SKAT kernel to improve detection power for rare variant associations. Through extensive simulations across diverse genetic architectures, we demonstrate that PrediXSKAT effectively controls the Type I error rate while substantially increasing statistical power compared to standard SKAT tests. Our results show that the method can uncover association signals that remain undetected in conventional approaches, particularly in scenarios characterized by sparse causal variants and mixed effect directions within a locus.

SKAT treats all carriers of rare variants in a gene equivalently regardless of the functional context in which those variants operate. PrediXSKAT modifies this by up-weighting rare variant similarity between individuals who also share similar predicted expression of that gene, and down-weighting similarity between individuals whose expression is divergent. We hypothesized that two individuals carrying rare variants in the same gene are more likely to share a phenotypically relevant perturbation when their baseline transcriptional output of that gene is also concordant. Our simulations demonstrate that simply including GReX as a covariate to capture the marginal effect of expression on the phenotype does not improve power consistently. The purpose of PrediXSKAT is not to account for a direct gene expression effect, but to modify the weights of the kernel to concentrate signal among individuals sharing both genetic and regulatory background.

Applying PrediXSKAT to whole-genome sequencing data from the Alzheimer’s Disease Sequencing Project (ADSP), we identify multiple genes associated with AD across three ancestry groups. PrediXSKAT recapitulates known AD susceptibility loci while highlighting previously unreported candidates supported by plausible biological mechanisms. Comparisons with SKAT analyses confirm that incorporating GReX information enhances sensitivity to biologically relevant signals while maintaining appropriate calibration. However, the improvement in significance from PrediXSKAT is most visible at moderate p-values, while genes with already-strong SKAT signal experience smaller shifts. This pattern is in part due to the low leverage of the *—* log_10_ *p* scale in that range. When a gene’s test statistic is already very large, even if PrediXSKAT increases the test statistic multiplicatively, the change in p-value does not reflect this shift.

The largest improvement is found in *PKHD1L1*, where the p-value is elevated from 6.60 *×* 10*^−^*^7^ to 1.91 *×* 10*^−^*^8^. This gene encodes fibrocystin-L, which is highly expressed on inner ear stereocilia and linked to hearing loss. Hearing impairment is one of the largest potentially modifiable risk factors for AD and related dementias across the life course [16]. *PKHD1L1* is also related to the *PKHD1* gene family, which is implicated in autosomal recessive polycystic kidney disease (ARPKD) (MIM: 263200), a primary ciliopathy [17]. Some clinical evidence links ARPKD with increased risk of dementia [18].

We observed modest genomic inflation under PrediXSKAT which warrants discussion. Genomic inflation factor *λ* is well-calibrated under the null for single-variant tests, but set-based tests like SKAT and PrediXSKAT aggregate multiple variants with varying allele frequencies, effect sizes, and LD patterns, which result in test statistics where the null distribution deviates from chi-squared distributions. We are also testing a much smaller set of genes relative to the number of variants tested in GWAS, so the median-based *λ* estimate is more sensitive to noise and outlier genes. The *λ* we observe may reflect residual correlation structure among aggregated variants within genes instead of systematic confounding. Our analyses of real-world data controlled for as many confounders as possible in the ADSP framework. For our users, inclusion of genomic relatedness matrix (or the sparse version) may be more effective in minimizing inflation.

Common variants and rare variants are often analyzed as separate genomic domains. In the past, the two variant classes would likely be from different sequencing technologies: common variants from genotyping arrays and rare variants from exome sequencing. As whole-genome sequencing has become more affordable, studying the interplay between them has become possible. We demonstrate a generalizable framework for a joint analysis of common regulatory and rare coding variants, based on empirical analysis of multi-ancestry cohorts with a complex disease phenotype.

Several limitations of the current study should be noted. First, our simulations were conducted exclusively using genotype data from the *ABCA7* locus, a known AD risk gene. While this provides realistic allele frequency distributions and linkage disequilibrium structures, *ABCA7* may not be representative of all regulatory architectures across the exome, and power estimates may be optimistic for smaller genes with fewer rare variants. The kernel parameterization was selected through preliminary analyses on the same dataset used for evaluation, which introduces the possibility of overfitting to *ABCA7*.

Future extensions of PrediXSKAT could address these limitations in several directions. Integration of multi-tissue GReX kernels, analogous to the multi-tissue TWAS frameworks, would allow the method to leverage correlated expression patterns across tissues. Incorporation of functional annotation weights alongside the GReX kernel, following the STAAR pipeline, could combine regulatory and variant-level biological priors in a unified test. More broadly, the framework of modulating rare variant kernels with continuous biological signals is not limited to gene expression, and analogous approaches using methylation, chromatin accessibility, or proteomics data could be developed within the same kernel construction paradigm.

## 5 Declarations & Acknowledgements

### Description of supplemental information

Supplemental information contains 3 figures.

## Acknowledgments

This work was supported by the following grants.

- NIA, RF1 AG070935 (Griswold, Bush)
- NIA, U19 AG074865 (Pericak-Vance, Vance, Tosto, Bush, Byrd, Reitz, Kunkle, Haines)
- NIA, U01 AG058654 (Haines, Bush, Martin, Farrer, Pericak-Vance)
- NIA, U54 AG052427 (Schellenberg, Wang)
- NIA, T32 GM135081 (Harris, Laframboise)
- NIEHS, T32 ES036169 (Ruden, Liu, Schumacher, Williams)

This work made use of the High Performance Computing Resource in the Core Facility for Advanced Research Computing at Case Western Reserve University.

## Data and code availability

Code for simulations with anonymized example data is available at: https://github.com/bushlab-genomics/predixskat

Data from the Alzheimer’s Disease Sequencing Project can be accessed through NIA-GADS (ng00067).

## Web Resources

- FaST-LMM: https://github.com/fastlmm/fastlmmclib
- PrediXcan: https://github.com/hakyimlab/MetaXcan
- GTEx (v8): https://predictdb.org/post/2021/07/21/gtex-v8-models-on-eqtl-and-sqtl/
- TWAS models from African American and Puerto Rican study cohorts: https://zenodo.org/records/7735723

## Author Contributions

Conceptualization, W.S.B. and Y.Y.; methodology, analyses and writing - original draft, Y.Y.; data and resources, E.R.M., L.A.F., M.A.P., J.M.V., J.L.H., W.S.B.; writing - review & editing, Y.Y. and W.S.B.; supervision, W.S.B.; all authors read and agreed to the final version of the manuscript.

## Alzheimer’s Disease Sequencing Project Acknowledgement Statement

The Alzheimer’s Disease Sequencing Project (ADSP) is comprised of two Alzheimer’s Disease (AD) genetics consortia and three National Human Genome Research Institute (NHGRI) funded Large Scale Sequencing and Analysis Centers (LSAC). The two AD genetics consortia are the Alzheimer’s Disease Genetics Consortium (ADGC) funded by NIA (U01 AG032984), and the Cohorts for Heart and Aging Research in Genomic Epidemiology (CHARGE) funded by NIA (R01 AG033193), the National Heart, Lung, and Blood Institute (NHLBI), other National Institute of Health (NIH) institutes and other foreign governmental and non-governmental organizations. The Discovery Phase analysis of sequence data is supported through UF1AG047133 (to Drs. Schellenberg, Farrer, Pericak-Vance, Mayeux, and Haines); U01AG049505 to Dr. Seshadri; U01AG049506 to Dr. Boerwinkle; U01AG049507 to Dr. Wijsman; and U01AG049508 to Dr. Goate and the Discovery Extension Phase analysis is supported through U01AG052411 to Dr. Goate, U01AG052410 to Dr. Pericak-Vance and U01 AG052409 to Drs. Seshadri and Fornage.

Sequencing for the Follow Up Study (FUS) is supported through U01AG057659 (to Drs. PericakVance, Mayeux, and Vardarajan) and U01AG062943 (to Drs. Pericak-Vance and Mayeux). Data generation and harmonization in the Follow-up Phase is supported by U54AG052427 (to Drs. Schellenberg and Wang). The FUS Phase analysis of sequence data is supported through U01AG058589 (to Drs. Destefano, Boerwinkle, De Jager, Fornage, Seshadri, and Wijsman), U01AG058654 (to Drs. Haines, Bush, Farrer, Martin, and Pericak-Vance), U01AG058635 (to Dr. Goate), RF1AG058066 (to Drs. Haines, Pericak-Vance, and Scott), RF1AG057519 (to Drs. Farrer and Jun), R01AG048927 (to Dr. Farrer), and RF1AG054074 (to Drs. Pericak-Vance and Beecham).

The ADGC cohorts include: Adult Changes in Thought (ACT) (U01 AG006781, U19 AG066567), the Alzheimer’s Disease Research Centers (ADRC) (P30 AG062429, P30 AG066468, P30 AG062421, P30 AG066509, P30 AG066514, P30 AG066530, P30 AG066507, P30 AG066444, P30 AG066518, P30 AG066512, P30 AG066462, P30 AG072979, P30 AG072972, P30 AG072976, P30 AG072975, P30 AG072978, P30 AG072977, P30 AG066519, P30 AG062677, P30 AG079280, P30 AG062422, P30 AG066511, P30 AG072946, P30 AG062715, P30 AG072973, P30 AG066506, P30 AG066508, P30 AG066515, P30 AG072947, P30 AG072931, P30 AG066546, P20 AG068024, P20 AG068053, P20 AG068077, P20 AG068082, P30 AG072958, P30 AG072959), the Chicago Health and Aging Project (CHAP) (R01 AG11101, RC4 AG039085, K23 AG030944), Indiana Memory and Aging Study (IMAS) (R01 AG019771), Indianapolis Ibadan (R01 AG009956, P30 AG010133), the Memory and Aging Project (MAP) ( R01 AG17917), Mayo Clinic (MAYO) (R01 AG032990, U01 AG046139, R01 NS080820, RF1 AG051504, P50 AG016574), Mayo Parkinson’s Disease controls (NS039764, NS071674, 5RC2HG005605), University of Miami (R01 AG027944, R01 AG028786, R01 AG019085, IIRG09133827, A2011048), the Multi-Institutional Research in Alzheimer’s Genetic Epidemiology Study (MIRAGE) (R01 AG09029, R01 AG025259), the National Centralized Repository for Alzheimer’s Disease and Related Dementias (NCRAD) (U24 AG021886), the National Institute on Aging Late Onset Alzheimer’s Disease Family Study (NIA-LOAD) (U24 AG056270), the Religious Orders Study (ROS) (P30 AG10161, R01 AG15819), the Texas Alzheimer’s Research and Care Consortium (TARCC) (funded by the Darrell K Royal Texas Alzheimer’s Initiative), Vanderbilt University/Case Western Reserve University (VAN/CWRU) (R01 AG019757, R01 AG021547, R01 AG027944, R01 AG028786, P01 NS026630, and Alzheimer’s Association), the Washington Heights-Inwood Columbia Aging Project (WHICAP) (RF1 AG054023), the University of Washington Families (VA Research Merit Grant, NIA: P50AG005136, R01AG041797, NINDS: R01NS069719), the Columbia University Hispanic Estudio Familiar de Influencia Genetica de Alzheimer (EFIGA) (RF1 AG015473), the University of Toronto (UT) (funded by Wellcome Trust, Medical Research Council, Canadian Institutes of Health Research), and Genetic Differences (GD) (R01 AG007584). The CHARGE cohorts are supported in part by National Heart, Lung, and Blood Institute (NHLBI) infrastructure grant HL105756 (Psaty), RC2HL102419 (Boerwinkle) and the neurology working group is supported by the National Institute on Aging (NIA) R01 grant AG033193.

The CHARGE cohorts participating in the ADSP include the following: Austrian Stroke Prevention Study (ASPS), ASPS-Family study, and the Prospective Dementia Registry-Austria (ASPS/PRODEM-Aus), the Atherosclerosis Risk in Communities (ARIC) Study, the Cardiovascular Health Study (CHS), the Erasmus Rucphen Family Study (ERF), the Framingham Heart Study (FHS), and the Rotterdam Study (RS). ASPS is funded by the Austrian Science Fond (FWF) grant number P20545-P05 and P13180 and the Medical University of Graz. The ASPS-Fam is funded by the Austrian Science Fund (FWF) project I904), the EU Joint Programme – Neurodegenerative Disease Research (JPND) in frame of the BRIDGET project (Austria, Ministry of Science) and the Medical University of Graz and the Steiermärkische Krankenanstalten Gesellschaft. PRODEM-Austria is supported by the Austrian Research Promotion agency (FFG) (Project No. 827462) and by the Austrian National Bank (Anniversary Fund, project 15435. ARIC research is carried out as a collaborative study supported by NHLBI contracts (HHSN268201100005C, HHSN268201100006C, HHSN268201100007C, HHSN268201100008C, HHSN268201100009C, HHSN268201100010C, HHSN268201100011C, and HHSN268201100012C). Neurocognitive data in ARIC is collected by U01 2U01HL096812, 2U01HL096814, 2U01HL096899, 2U01HL096902, 2U01HL096917 from the NIH (NHLBI, NINDS, NIA and NIDCD), and with previous brain MRI examinations funded by R01-HL70825 from the NHLBI. CHS research was supported by contracts HHSN268201200036C, HHSN268200800007C, N01HC55222, N01HC85079, N01HC85080, N01HC85081, N01HC85082, N01HC85083, N01HC85086, and grants U01HL080295 and U01HL130114 from the NHLBI with additional contribution from the National Institute of Neurological Disorders and Stroke (NINDS). Additional support was provided by R01AG023629, R01AG15928, and R01AG20098 from the NIA. FHS research is supported by NHLBI contracts N01-HC-25195 and HHSN268201500001I. This study was also supported by additional grants from the NIA (R01s AG054076, AG049607 and AG033040 and NINDS (R01 NS017950). The ERF study as a part of EUROSPAN (European Special Populations Research Network) was supported by European Commission FP6 STRP grant number 018947 (LSHG-CT-2006-01947) and also received funding from the European Community’s Seventh Framework Programme (FP7/2007-2013)/grant agreement HEALTH-F4-2007-201413 by the European Commission under the programme “Quality of Life and Management of the Living Resources” of 5th Framework Programme (no. QLG2-CT-2002-01254). High-throughput analysis of the ERF data was supported by a joint grant from the Netherlands Organization for Scientific Research and the Russian Foundation for Basic Research (NWO-RFBR 047.017.043). The Rotterdam Study is funded by Erasmus Medical Center and Erasmus University, Rotterdam, the Netherlands Organization for Health Research and Development (ZonMw), the Research Institute for Diseases in the Elderly (RIDE), the Ministry of Education, Culture and Science, the Ministry for Health, Welfare and Sports, the European Commission (DG XII), and the municipality of Rotterdam. Genetic data sets are also supported by the Netherlands Organization of Scientific Research NWO Investments (175.010.2005.011, 911-03-012), the Genetic Laboratory of the Department of Internal Medicine, Erasmus MC, the Research Institute for Diseases in the Elderly (014-93-015; RIDE2), and the Netherlands Genomics Initiative (NGI)/Netherlands Organization for Scientific Research (NWO) Netherlands Consortium for Healthy Aging (NCHA), project 050-060-810. All studies are grateful to their participants, faculty and staff. The content of these manuscripts is solely the responsibility of the authors and does not necessarily represent the official views of the National Institutes of Health or the U.S. Department of Health and Human Services.

The FUS cohorts include: the Alzheimer’s Disease Research Centers (ADRC) (P30 AG062429, P30 AG066468, P30 AG062421, P30 AG066509, P30 AG066514, P30 AG066530, P30 AG066507, P30 AG066444, P30 AG066518, P30 AG066512, P30 AG066462, P30 AG072979, P30 AG072972, P30 AG072976, P30 AG072975, P30 AG072978, P30 AG072977, P30 AG066519, P30 AG062677, P30 AG079280, P30 AG062422, P30 AG066511, P30 AG072946, P30 AG062715, P30 AG072973, P30 AG066506, P30 AG066508, P30 AG066515, P30 AG072947, P30 AG072931, P30 AG066546, P20 AG068024, P20 AG068053, P20 AG068077, P20 AG068082, P30 AG072958, P30 AG072959), Alzheimer’s Disease Neuroimaging Initiative (ADNI) (U19AG024904), Amish Protective Variant Study (RF1AG058066), Cache County Study (R01AG11380, R01AG031272, R01AG21136, RF1AG054052), Case Western Reserve University Brain Bank (CWRUBB) (P50AG008012), Case Western Reserve University Rapid Decline (CWRURD) (RF1AG058267, NU38CK000480), CubanAmerican Alzheimer’s Disease Initiative (CuAADI) (3U01AG052410), Estudio Familiar de Influencia Genetica en Alzheimer (EFIGA) (5R37AG015473, RF1AG015473, R56AG051876), Genetic and Environmental Risk Factors for Alzheimer Disease Among African Americans Study (GenerAAtions) (2R01AG09029, R01AG025259, 2R01AG048927), Gwangju Alzheimer and Related Dementias Study (GARD) (U01AG062602), Hillblom Aging Network (2014-A-004-NET, R01AG032289, R01AG048234), Hussman Institute for Human Genomics Brain Bank (HIHGBB) (R01AG027944, Alzheimer’s Association “Identification of Rare Variants in Alzheimer Disease”), Ibadan Study of Aging (IBADAN) (5R01AG009956), Longevity Genes Project (LGP) and LonGenity (R01AG042188, R01AG044829, R01AG046949, R01AG057909, R01AG061155, P30AG038072), Mexican Health and Aging Study (MHAS) (R01AG018016), Multi-Institutional Research in Alzheimer’s Genetic Epidemiology (MIRAGE) (2R01AG09029, R01AG025259, 2R01AG048927), Northern Manhattan Study (NOMAS) (R01NS29993), Peru Alzheimer’s Disease Initiative (PeADI) (RF1AG054074), Puerto Rican 1066 (PR1066) (Wellcome Trust (GR066133/GR080002), European Research Council (340755)), Puerto Rican Alzheimer Disease Initiative (PRADI) (RF1AG054074), Reasons for Geographic and Racial Differences in Stroke (REGARDS) (U01NS041588), Research in African American Alzheimer Disease Initiative (REAAADI) (U01AG052410), the Religious Orders Study (ROS) (P30 AG10161, P30 AG72975, R01 AG15819, R01 AG42210), the RUSH Memory and Aging Project (MAP) (R01 AG017917, R01 AG42210Stanford Extreme Phenotypes in AD (R01AG060747), University of Miami Brain Endowment Bank (MBB), University of Miami/Case Western/North Carolina A&T African American (UM/CASE/NCAT) (U01AG052410, R01AG028786), Wisconsin Registry for Alzheimer’s Prevention (WRAP) (R01AG027161 and R01AG054047), Mexico-Southern California Autosomal Dominant Alzheimer’s Disease Consortium (R01AG069013), Center for Cognitive Neuroscience and Aging (R01AG047649), and the A4 Study (R01AG063689, U19AG010483 and U24AG057437).

The four LSACs are: the Human Genome Sequencing Center at the Baylor College of Medicine (U54 HG003273), the Broad Institute Genome Center (U54HG003067), The American Genome Center at the Uniformed Services University of the Health Sciences (U01AG057659), and the Washington University Genome Institute (U54HG003079). Genotyping and sequencing for the ADSP FUS is also conducted at John P. Hussman Institute for Human Genomics (HIHG) Center for Genome Technology (CGT).

Biological samples and associated phenotypic data used in primary data analyses were stored at Study Investigators institutions, and at the National Centralized Repository for Alzheimer’s Disease and Related Dementias (NCRAD, U24AG021886) at Indiana University funded by NIA. Associated Phenotypic Data used in primary and secondary data analyses were provided by Study Investigators, the NIA funded Alzheimer’s Disease Centers (ADCs), and the National Alzheimer’s Coordinating Center (NACC, U24AG072122) and the National Institute on Aging Genetics of Alzheimer’s Disease Data Storage Site (NIAGADS, U24AG041689) at the University of Pennsylvania, funded by NIA. Harmonized phenotypes were provided by the ADSP Phenotype Harmonization Consortium (ADSP-PHC), funded by NIA (U24 AG074855, U01 AG068057 and R01 AG059716) and Ultrascale Machine Learning to Empower Discovery in Alzheimer’s Disease Biobanks (AI4AD, U01 AG068057). This research was supported in part by the Intramural Research Program of the National Institutes of health, National Library of Medicine. Contributors to the Genetic Analysis Data included Study Investigators on projects that were individually funded by NIA, and other NIH institutes, and by private U.S. organizations, or foreign governmental or nongovernmental organizations.

The ADSP Phenotype Harmonization Consortium (ADSP-PHC) is funded by NIA (U24 AG074855, U01 AG068057 and R01 AG059716). The harmonized cohorts within the ADSP-PHC include: the Anti-Amyloid Treatment in Asymptomatic Alzheimer’s study (A4 Study), a secondary prevention trial in preclinical Alzheimer’s disease, aiming to slow cognitive decline associated with brain amyloid accumulation in clinically normal older individuals. The A4 Study is funded by a public-private-philanthropic partnership, including funding from the National Institutes of Health-National Institute on Aging, Eli Lilly and Company, Alzheimer’s Association, Accelerating Medicines Partnership, GHR Foundation, an anonymous foundation and additional private donors, with in-kind support from Avid and Cogstate. The companion observational Longitudinal Evaluation of Amyloid Risk and Neurodegeneration (LEARN) Study is funded by the Alzheimer’s Association and GHR Foundation. The A4 and A4-LEARN Studies are led by Dr. Reisa Sperling at Brigham and Women’s Hospital, Harvard Medical School and Dr. Paul Aisen at the Alzheimer’s Therapeutic Research Institute (ATRI), University of Southern California. The A4 and LEARN Studies are coordinated by ATRI at the University of Southern California, and the data are made available through the Laboratory for Neuro Imaging at the University of Southern California. The participants screening for the A4 Study provided permission to share their de-identified data in order to advance the quest to find a successful treatment for Alzheimer’s disease. We would like to acknowledge the dedication of all the participants, the site personnel, and all of the partnership team members who continue to make the A4 and LEARN Studies possible. The complete A4 Study Team list is available on: a4study.org/a4-study-team.; the Adult Changes in Thought study (ACT), U01 AG006781, U19 AG066567; Alzheimer’s Disease Neuroimaging Initiative (ADNI): Data collection and sharing for this project was funded by the Alzheimer’s Disease Neuroimaging Initiative (ADNI) (National Institutes of Health Grant U01 AG024904) and DOD ADNI (Department of Defense award number W81XWH-12-2-0012). ADNI is funded by the National Institute on Aging, the National Institute of Biomedical Imaging and Bioengineering, and through generous contributions from the following: AbbVie, Alzheimer’s Association; Alzheimer’s Drug Discovery Foundation; Araclon Biotech; BioClinica, Inc.; Biogen; Bristol-Myers Squibb Company; CereSpir, Inc.; Cogstate; Eisai Inc.; Elan Pharmaceuticals, Inc.; Eli Lilly and Company; EuroImmun; F. Hoffmann-La Roche Ltd and its affiliated company Genentech, Inc.; Fujirebio; GE Healthcare; IXICO Ltd.; Janssen Alzheimer Immunotherapy Research & Development, LLC.; Johnson & Johnson Pharmaceutical Research & Development LLC.; Lumosity; Lundbeck; Merck & Co., Inc.; Meso Scale Diagnostics, LLC.; NeuroRx Research; Neurotrack Technologies; Novartis Pharmaceuticals Corporation; Pfizer Inc.; Piramal Imaging; Servier; Takeda Pharmaceutical Company; and Transition Therapeutics. The Canadian Institutes of Health Research is providing funds to support ADNI clinical sites in Canada. Private sector contributions are facilitated by the Foundation for the National Institutes of Health (www.fnih.org). The grantee organization is the Northern California Institute for Research and Education, and the study is coordinated by the Alzheimer’s Therapeutic Research Institute at the University of Southern California. ADNI data are disseminated by the Laboratory for Neuro Imaging at the University of Southern California; Brain tissue and additional biological samples were collected from individuals who were assessed and followed in the Case Western ADC which was funded by NIA P50AG008012; Estudio Familiar de Influencia Genetica en Alzheimer (EFIGA): 5R37AG015473, RF1AG015473, R56AG051876; the Health & Aging Brain Study – Health Disparities (HABS-HD), supported by the National Institute on Aging of the National Institutes of Health under Award Numbers R01AG054073, R01AG058533, R01AG070862, P41EB015922, and U19AG078109; brain tissues were obtained from the Brain Bank at the John P. Hussman Institute for Human Genomics at the University of Miami Miller School of Medicine, with funding provided by Louis D. Scientific Prize from the Institut de France; the Indiana Alzheimer’s Disease Research Center (IADRC) supported by National Institutes of Health (NIH) grant P30AG072976; the Korean Brain Aging Study for the Early Diagnosis and Prediction of Alzheimer’s disease (KBASE), which was supported by a grant from Ministry of Science, ICT and Future Planning (Grant No: NRF-2014M3C7A1046042); Memory & Aging Project at Knight Alzheimer’s Disease Research Center (MAP at Knight ADRC): The Memory and Aging Project at the Knight-ADRC (Knight-ADRC). This work was supported by the National Institutes of Health (NIH) grants R01AG064614, R01AG044546, RF1AG053303, RF1AG058501, U01AG058922 and R01AG064877 to Carlos Cruchaga. The recruitment and clinical characterization of research participants at Washington University was supported by NIH grants P30AG066444, P01AG03991, and P01AG026276. Data collection and sharing for this project was supported by NIH grants RF1AG054080, P30AG066462, R01AG064614 and U01AG052410. We thank the contributors who collected samples used in this study, as well as patients and their families, whose help and participation made this work possible. This work was supported by access to equipment made possible by the Hope Center for Neurological Disorders, the Neurogenomics and Informatics Center (NGI: https://neurogenomics.wustl.edu/) and the Departments of Neurology and Psychiatry at Washington University School of Medicine; the University of Miami Brain Endowment Bank (Miami Brain Bank); National Alzheimer’s Coordinating Center (NACC): The NACC database is funded by NIA/NIH Grant U24 AG072122. SCAN is a multi-institutional project that was funded as a U24 grant (AG067418) by the National Institute on Aging in May 2020. Data collected by SCAN and shared by NACC are contributed by the NIA-funded ADRCs as follows: P30 AG062429 (PI James Brewer, MD, PhD), P30 AG066468 (PI Oscar Lopez, MD), P30 AG062421 (PI Bradley Hyman, MD, PhD), P30 AG066509 (PI Thomas Grabowski, MD), P30 AG066514 (PI Mary Sano, PhD), P30 AG066530 (PI Helena Chui, MD), P30 AG066507 (PI Marilyn Albert, PhD), P30 AG066444 (PI John Morris, MD), P30 AG066518 (PI Jeffrey Kaye, MD), P30 AG066512 (PI Thomas Wisniewski, MD), P30 AG066462 (PI Scott Small, MD), P30 AG072979 (PI David Wolk, MD), P30 AG072972 (PI Charles DeCarli, MD), P30 AG072976 (PI Andrew Saykin, PsyD), P30 AG072975 (PI David Bennett, MD), P30 AG072978 (PI Neil Kowall, MD), P30 AG072977 (PI Robert Vassar, PhD), P30 AG066519 (PI Frank LaFerla, PhD), P30 AG062677 (PI Ronald Petersen, MD, PhD), P30 AG079280 (PI Eric Reiman, MD), P30 AG062422 (PI Gil Rabinovici, MD), P30 AG066511 (PI Allan Levey, MD, PhD), P30 AG072946 (PI Linda Van Eldik, PhD), P30 AG062715 (PI Sanjay Asthana, MD, FRCP), P30 AG072973 (PI Russell Swerdlow, MD), P30 AG066506 (PI Todd Golde, MD, PhD), P30 AG066508 (PI Stephen Strittmatter, MD, PhD), P30 AG066515 (PI Victor Henderson, MD, MS), P30 AG072947 (PI Suzanne Craft, PhD), P30 AG072931 (PI Henry Paulson, MD, PhD), P30 AG066546 (PI Sudha Seshadri, MD), P20 AG068024 (PI Erik Roberson, MD, PhD), P20 AG068053 (PI Justin Miller, PhD), P20 AG068077 (PI Gary Rosenberg, MD), P20 AG068082 (PI Angela Jefferson, PhD), P30 AG072958 (PI Heather Whitson, MD), P30 AG072959 (PI James Leverenz, MD); National Institute on Aging Alzheimer’s Disease Family Based Study (NIA-AD FBS): U24 AG056270; Religious Orders Study (ROS): P30AG10161, R01AG15819, R01AG42210; Memory and Aging Project (MAP - Rush): R01AG017917, R01AG42210; Minority Aging Research Study (MARS): R01AG22018, R01AG42210; the Texas Alzheimer’s Research and Care Consortium (TARCC), funded by the Darrell K Royal Texas Alzheimer’s Initiative, directed by the Texas Council on Alzheimer’s Disease and Related Disorders; Washington Heights/Inwood Columbia Aging Project (WHICAP): RF1 AG054023; and Wisconsin Registry for Alzheimer’s Prevention (WRAP): R01AG027161 and R01AG054047. Additional acknowledgments include the National Institute on Aging Genetics of Alzheimer’s Disease Data Storage Site (NIAGADS, U24AG041689) at the University of Pennsylvania, funded by NIA.

## 8 Supplementary Information

**Fig. S1.**
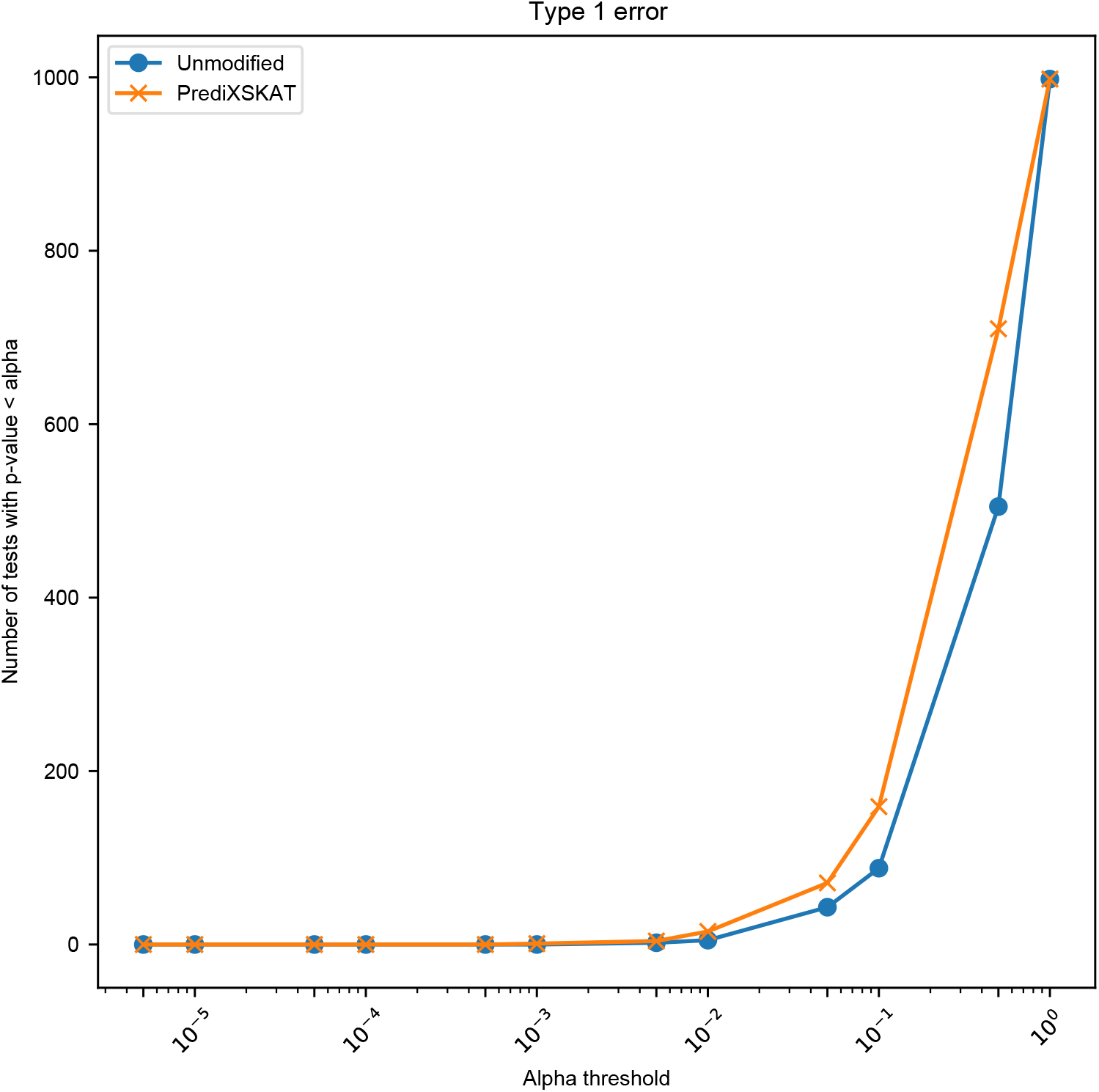
Type I error of PrediXSKAT and unmodified SKAT across a range of. *α* **thresholds.** For each *α* threshold (x-axis, log scale), the y-axis shows the number of null-model simulation replicates with p *< α*.

**Fig. S2.**
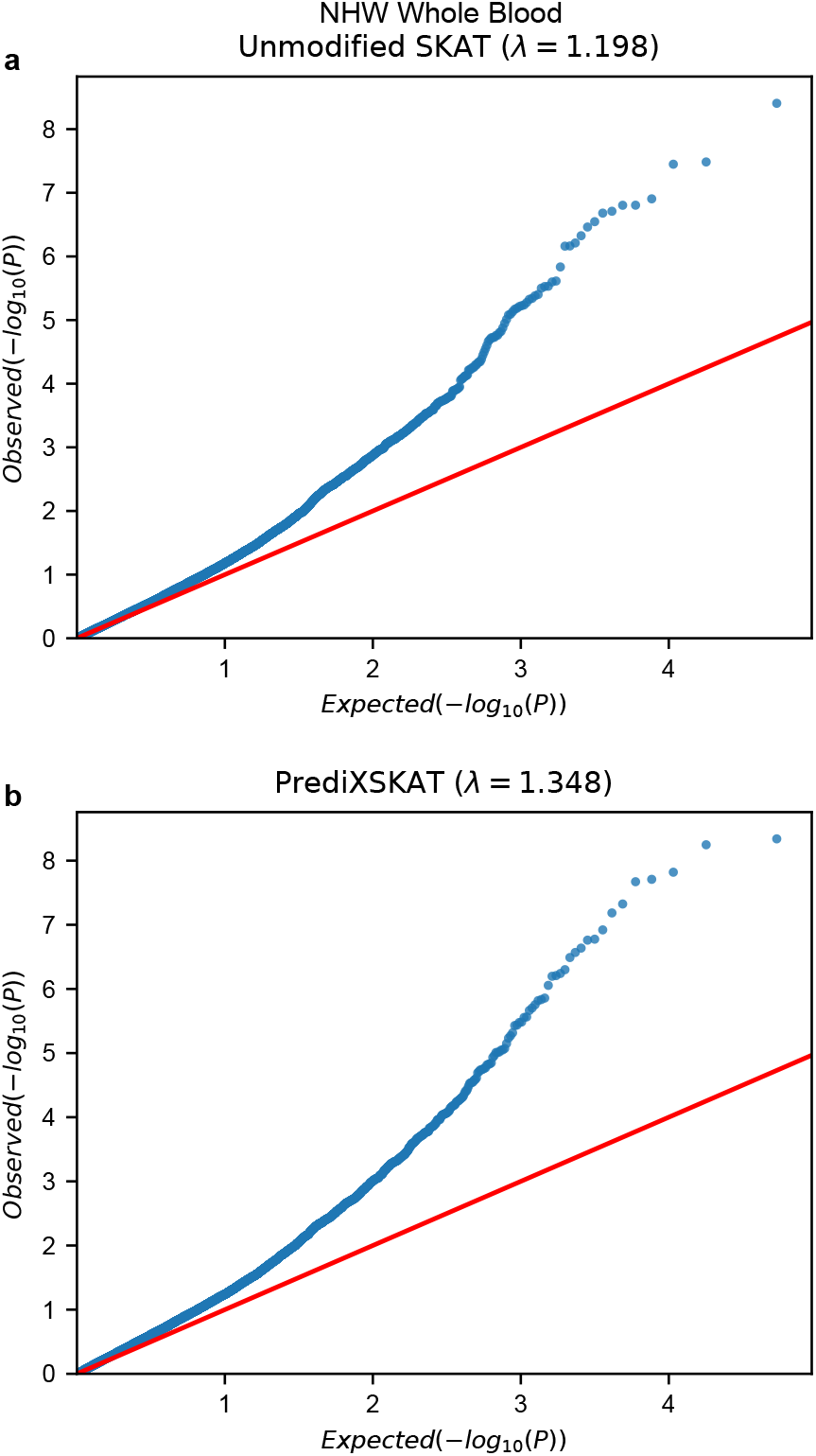
Quantile–quantile plots of gene-based rare-variant test p-values for the NHW whole-blood analysis (same data as. **Figure 8)** (a) Unmodified SKAT (genomic inflation factor *λ* = 1.198). (b) PrediXSKAT (*λ* = 1.348). Each point represents one gene.

**Fig. S3.**
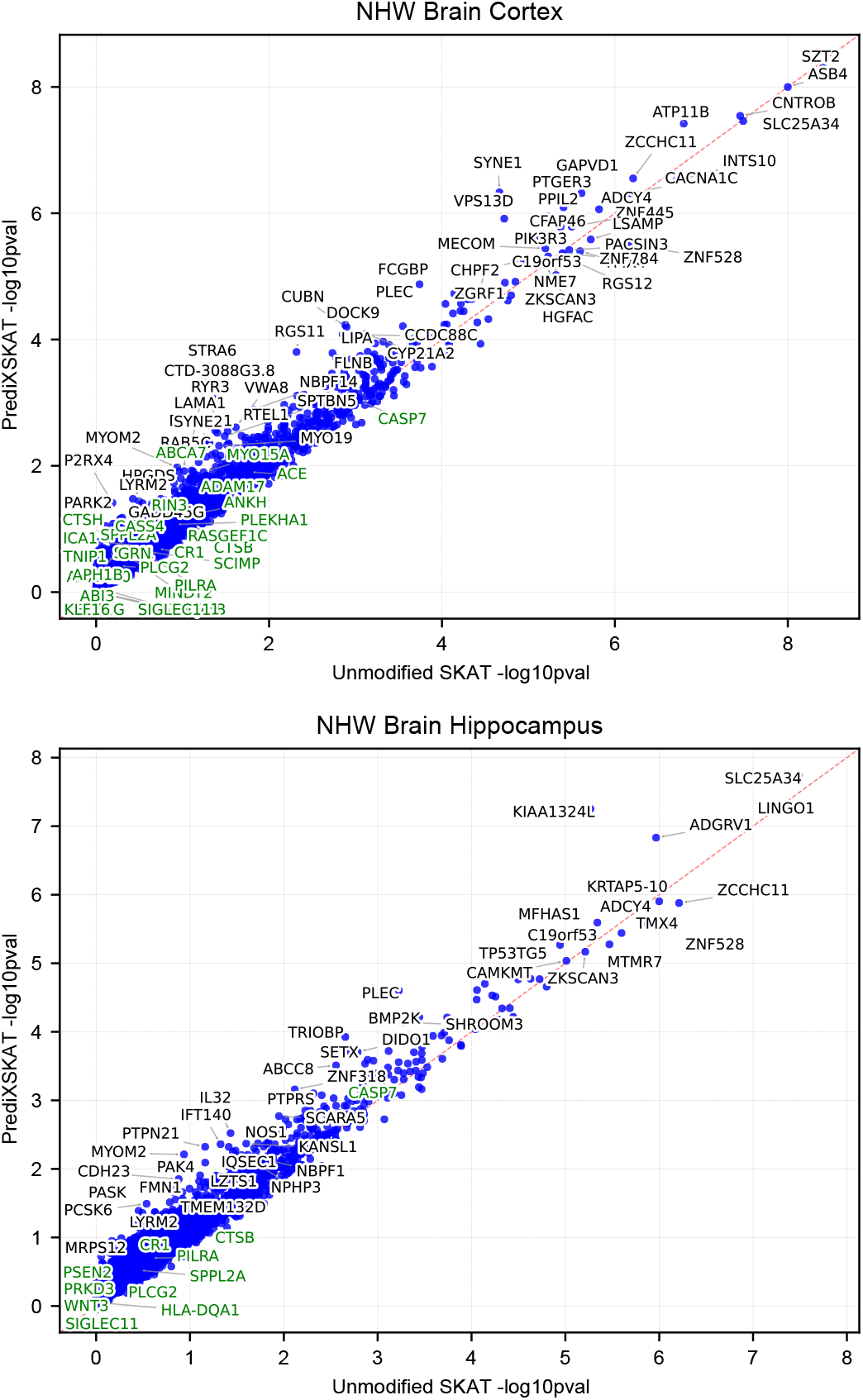
PrediXSKAT versus unmodified SKAT results using GReX modeled from brain tissue, NHW subgroup. (a) Brain cortex GReX model. (b) Hippocampus GReX model. Plotted as in Figure 8.

